# A membrane-associated conveyor belt controls the rotational direction of the bacterial type 9 secretion system

**DOI:** 10.1101/2024.09.23.614571

**Authors:** Abhishek Trivedi, Jacob A. Miratsky, Emma C. Henderson, Abhishek Singharoy, Abhishek Shrivastava

## Abstract

Many bacteria utilize the type 9 secretion system (T9SS) for gliding motility, surface colonization, and pathogenesis. This dual-function motor supports both gliding motility and protein secretion, where rotation of the T9SS plays a central role. Fueled by the energy of the stored proton motive force and transmitted through the torque of membrane-anchored stator units, the rotary T9SS propels an adhesin-coated conveyor belt along the bacterial outer membrane like a molecular snowmobile, thereby enabling gliding motion. However, the mechanisms controlling the rotational direction and gliding motility of T9SS remain elusive. Shedding light on this mechanism, we find that in the gliding bacterium *Flavobacterium johnsoniae*, deletion of the C-terminus of a conveyor belt protein GldJ controls, and in fact, reverses the rotational direction of T9SS from counterclockwise to clockwise thus suggesting that the interface between the conveyor belt protein GldJ and the T9SS ring protein GldK plays an important role in controlling the directionality of T9SS. Combined with MD simulation of the T9SS stator units GldLM, we suggest a ‘tri-component gearset’ model where the conveyor belt controls the rotational direction of its driver, the T9SS, thus providing adaptive sensory feedback to influence the motility of the gliding bacterium.

## INTRODUCTION

To date, three classes of ion-driven biological rotary motors have been identified: the type 9 secretion system (T9SS), ATP synthase, and the bacterial flagellar motor. The T9SS, discovered recently, is utilized by the Bacteroidetes-chlorobi-fibrobacteres superphylum bacteria. Over 200,000 distinct proteins secreted by T9SS have been cataloged on the InterPro^1^, showcasing a remarkable diversity. These proteins span a wide range of functions and include motility adhesins^2^, virulent proteases^3,4^, and enzymes that break down complex polysaccharides^5,6^. Changes in Bacteroidetes abundances in human microbiota correlate with several diseases. The T9SS of a gut Bacteroidetes, *Paraprevotella clara*, plays a crucial role in defending against viral infections^7^. In contrast, in the case of oral microbiota, the inflammatory response amplification induced by enzymes secreted by T9SS of *Porphyromonas gingivalis* is associated with the onset of periodontitis and Alzheimer’s disease^8,9^. Furthermore, the T9SS is required for the development of virulent biofilms^10–12^, which are harmful to a variety of hosts including humans^9^, birds^13^, and fish^14,15^.

T9SS also enables the gliding motility of Bacteroidetes over human and animal tissues, fish scales, sediments, and plant roots^16–18^. *Flavobacterium johnsoniae*, found in the soybean rhizosphere^19^, is a model organism for investigation of both the macromolecular mechanics of T9SS and the gliding motility of Bacteroidetes. T9SS rotates in place around a stationary axis and its rotational motion is driven by a proton motive force^3^. Following measurements of fluorescently labeled T9SS, tracking of motility adhesins on the conveyor belt, and tethered cell analysis, it has been suggested that similar to a macromolecular rack and pinion assembly, T9SS propels a polymeric cell-surface conveyor belt over the cell body, thus mobilizing a molecular snowmobile that enables the gliding motility of Bacteroidetes^20^ (Fig. 1A). Generally, T9SS stands out as a multifunctional apparatus - it is known to provide competitive advantage to a cell by converting the energy of ionic flow into rotational motion, with the generated rotational energy being utilized to enable either secretion processes, or direct mechanical coupling with a conveyor belt, thus facilitating cellular locomotion. Although gliding motility represents a distinctive example of cellular evolution, essentially nothing is known about molecular mechanisms regulating rotational direction or the mechanical linkage between T9SS and the conveyor belt. In *F. johnsoniae*, known for its robust gliding, the rotation of the GldKN ring propels the linear motion of a cell-surface adhesin, SprB, along a conveyor belt made of the protein GldJ^21–23^. The direct surface interaction between SprB and an external substratum enables gliding motility^21^. Intriguingly, GldJ is missing from T9SS in non-motile bacteria which utilize T9SS exclusively for protein secretion and pathogenesis. For instance, *P. gingivalis* is known to possess a large GldKN (PorKN) ring secretes virulent gingipain proteases via T9SS^8^.

**Figure 1.**
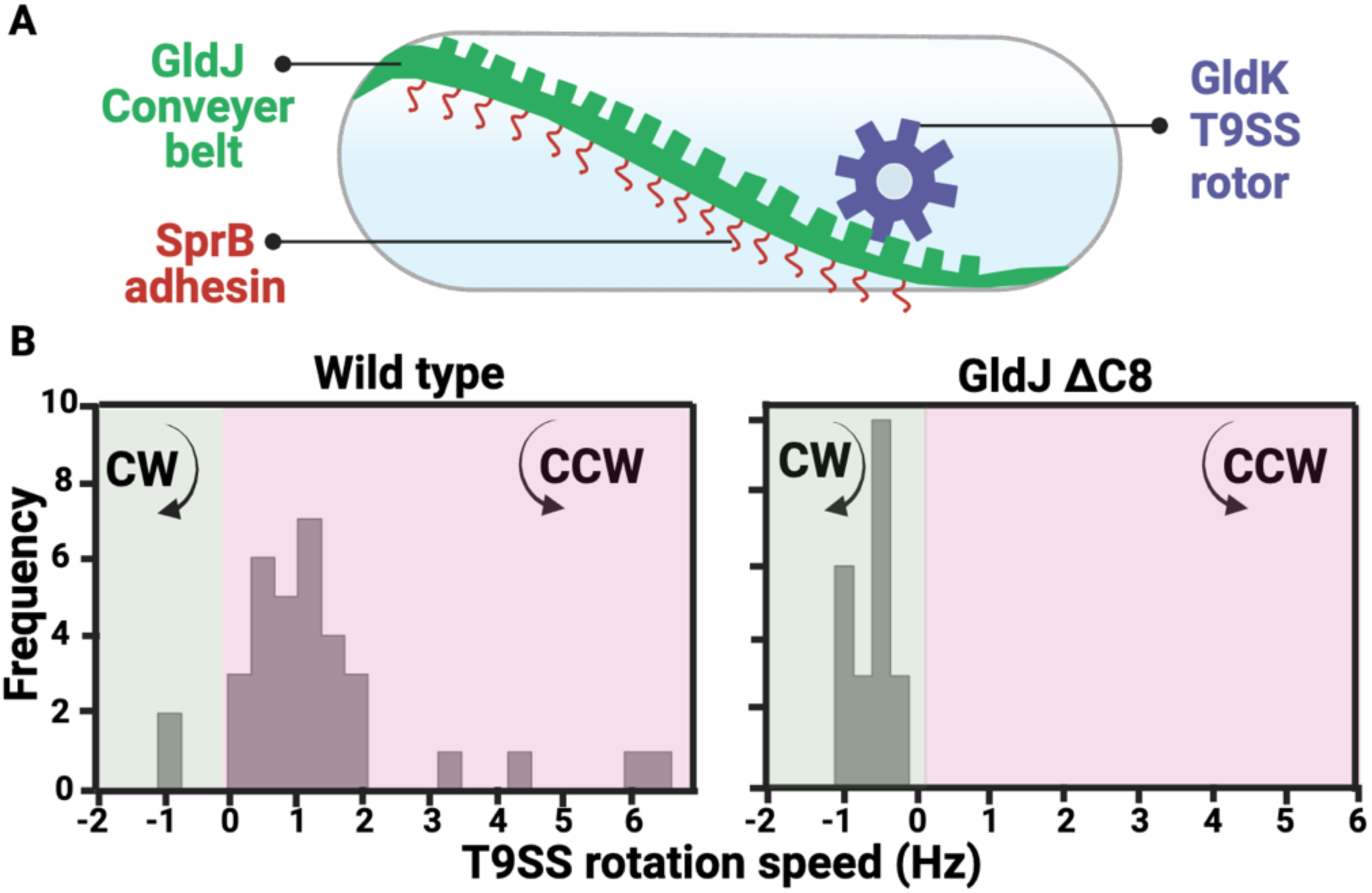
The C-terminal region of a conveyor belt protein, GldJ controls the rotational direction of T9SS. **(A)** A cartoon showing the current rack and pinion model where the GldK ring of rotary T9SS (pinion) drives the GldJ conveyor belt (rack). **(B)** Frequency distribution of rotational speed shows that T9SS of wild-type cells (n= 33) rotate primarily in the counterclockwise (CCW) direction whereas T9SS of cells lacking the C-terminal region of GldJ (GldJ ΔC8, n= 21) rotate in the clockwise (CW) direction.

T9SS consists of around twenty unique proteins working together to enable both gliding motility and protein secretion^24^. These building-block proteins do not exhibit sequence homology with the extensively studied bacterial flagellar motor that enables swimming motility of other bacteria. The mechanical motion in T9SS is enabled by proton conducting GldLM stator units exhibiting 5:2 structural stoichiometry. Predominantly, GldL is cytoplasmic with transmembrane helices forming a pentameric cage which encloses a periplasmic GldM dimer. GldL localizes on the rotational axis of tethered cells, and in gliding cells it remains stationary^25,26^. The GldM (PorM) periplasmic arm has 18-fold symmetry, and it interacts with the outer-membrane embedded GldKN (PorKN) ring^27^. Macromolecular structure and 5:2 stoichiometry of GldLM is both reminiscent of the flagellar MotAB proteins, suggesting somewhat similar principles in their evolution. T9SS and bacterial flagellar motor both use ion channels to spin a ring but the location of the ring varies. Specifically, in bacterial flagellar motor proton-conducting MotAB stator units drive rotational motion of a cytoplasmic ring (C-ring), and the rotational bias is controlled by a cytoplasmic protein sensory transduction protein, CheY-P, to the C-ring^28,29^. In contrast, in T9SS, the GldLM-driven rotation of GldKN ring occurs in the outer membrane. Extending the logic that allosteric changes in its C-ring determine the rotational direction in bacterial flagellar motor, it could be hypothesized that some protein binding to the GldKN ring might regulate the T9SS rotational bias. This suggests the presence of an outer membrane-associated sensory transducer, but any evidence is currently absent. The gliding motility of Bacteroidetes showcases seamless coordination of multiple macromolecular units that drive directed cellular motion, yet its actual macromolecular mechanisms have long remained enigmatic^30,31^. Here, we find that an unlikely candidate, namely, the cell-surface conveyor belt protein GldJ controls the rotational bias in T9SS. Deletion of 8 amino acids from the C-terminus of the conveyor belt protein GldJ (GldJ ΔC8) switches T9SS from CCW to CW direction. Further suppressor mutagenesis revealed mutations in the T9SS ring protein GldK which partially complemented the loss of GldJ’s C-terminal region, suggesting conservation of interaction energy between the T9SS ring protein GldK and the conveyor belt protein GldJ. The T9SS motor is driven by direct interaction of individual GldM proteins within GldLM stator units with the GldKN ring^25^. Unraveling the dynamics of the stator, our steered and Bayesian MD simulations suggest a CW rotational bias for GldM stator hence the directionality of T9SS is controlled at the interface of the GldK-GldJ interface. Combined, our results here suggest that GldLM stator units, GldKN ring, and GldJ conveyor belt constitute a tri-component gearset driving the gliding motility in Bacteroidetes. Namely, the stator complex harnesses proton motive force to propel the GldKN ring which, in its turn, facilitates and controls the mechanical motion of a closed GldJ conveyor belt. Our model suggests a unique molecular mechanism which alters the directionality of cellular motility, whereby a component of the cell-surface conveyor belt (GldJ) enables a regulatory feedback loop influencing the rotational bias of its associated gear (GldKN ring) and thereby altering the directionality of the propulsive motion of the conveyor belt.

## RESULTS

### Conveyer-belt protein GldJ controls directionality of T9SS motor

It has been reported that, in wild-type *F. johnsoniae*, 90% of T9SS units rotate CCW, whereas the remaining 10% rotate CW^3^. We gathered additional data and quantified rotational preferences of tethered T9SS in sheared wild-type cells. In agreement with earlier findings, our data confirms that T9SS is capable of rotating bi-directionally with a bias toward CCW direction (Fig. 1B, Supplementary video 1). T9SS motors appear to drive the cell-surface GldJ protein of the conveyor belt^32^ (Fig. 1A). Deletion of GldJ destabilizes GldK and cellular levels of GldK are significantly reduced^33^. Yet, when 8 specific amino acids on the C-terminus of GldJ are deleted (GldJ ΔC8), GldK levels are maintained. However, despite near-wild-type levels of GldK, GldJ ΔC8 cells do not exhibit swarming motility on agar^33^. These observations suggest that GldJ might somehow influence the rotation or assembly of T9SS. To test this conjecture, GldJ ΔC8 cells were sheared and tethered to a glass surface. Surprisingly, our experimental findings revealed a distinct change in T9SS rotational directionality as tethered T9SS motors rotated exclusively CW in GldJ ΔC8 cells (Fig. 1B, Supplementary video 2). In wild-type cells, T9SS exhibited an average rotational speed of 1.3 Hz (CCW) whereas in GldJ ΔC8 cells the rotational speed was about 0.6 Hz (CW, Fig. 1B). The above results indicate that the 8 C-terminal amino acids of GldJ play a crucial role in regulating both the speed and directionality of the T9SS motor. In the absence of those amino acids of the conveyor belt protein GldJ, T9SS rotates exclusively CW.

### Suppressor mutagenesis revealed that point mutations in GldK in GldJ ΔC8 cells restored swarming

Wild-type *F. johnsoniae* are known to robustly swarm on PY2 agar, whereas GldJ ΔC8 cells do not swarm on PY2 agar^33^. Suppressor mutagenesis of GldJ ΔC8 cells yielded strains that were capable of swarming on agar (Fig. 2A, Fig. S1). Three individual suppressor strains were isolated and their whole genome was sequenced. All the three turned out to be point mutations in the T9SS ring protein GldK, with the mutations occurring at the R73S, N307S, and M77L sites of the strains, respectively, further supporting the fact that GldJ and GldK interact closely with each other^23^ .

**Figure 2.**
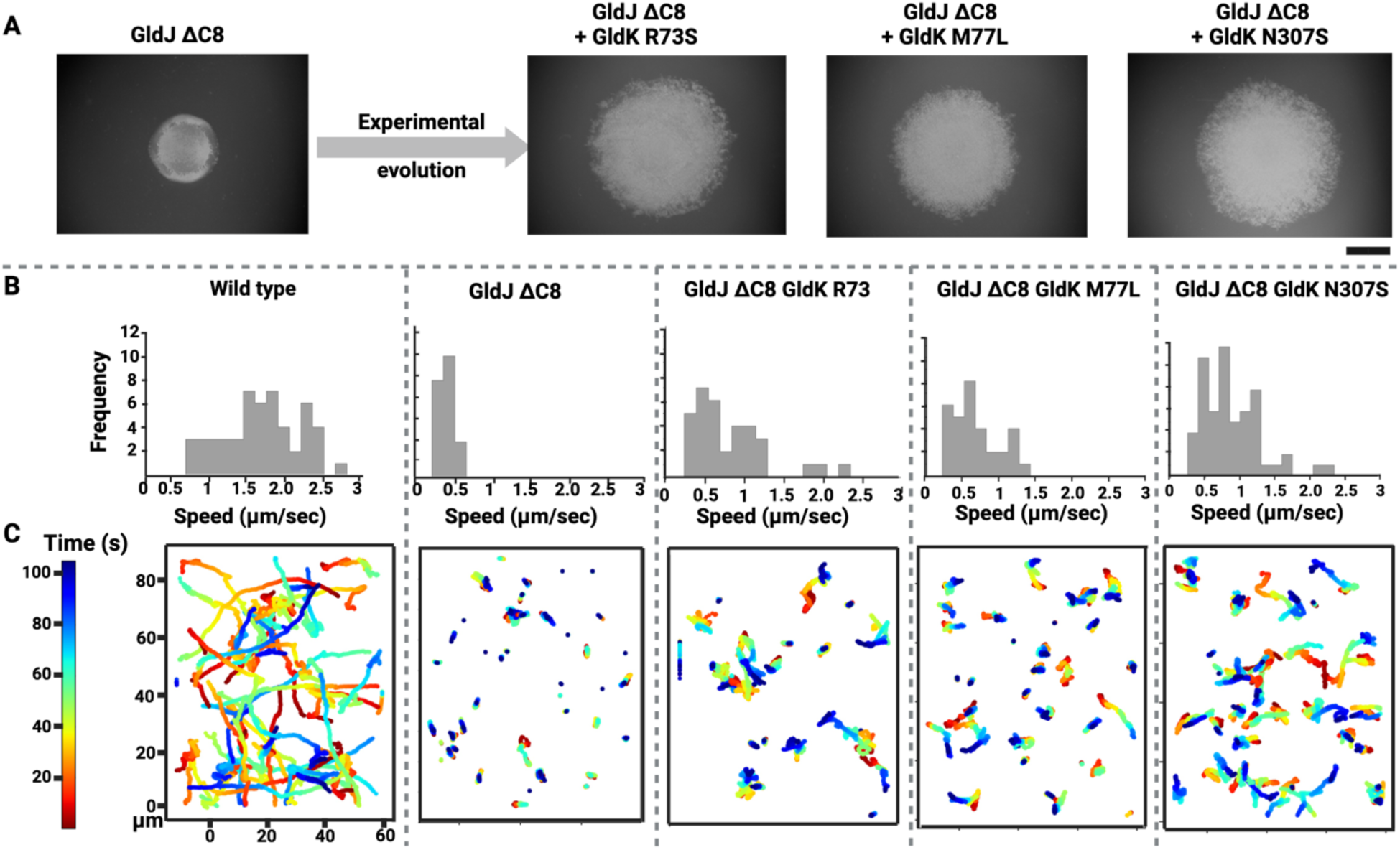
Experimental evolution restored gliding motility in cells that lack the C-terminal region of GldJ. **(A)** Cells lacking the C-terminal region of GldJ (GldJ ΔC8) do not swarm on agar. Experimental evolution of the GldJ ΔC8 strain resulted in partial restoration of swarming. All spontaneous mutations in the evolved strains were in the T9SS ring protein GldK. Scale bar = 6 mm. **(B)** Frequency distribution of the speed with which wild-type and mutant cells glide over a glass surface. **(C)** Trajectories of wild-type and mutant cells gliding over a glass surface (n=30).

Single-cell motility over a glass surface of the wild-type, GldJ ΔC8 strain, and the GldK suppressor strains was recorded, and individual trajectories of cellular motion were plotted. Wild-type cells exhibited long runs followed by relatively quick turning events characterized by either back-and-forth cellular motion or rare head-on flips, whereas GldJ ΔC8 cells mostly exhibited minimal back-and-forth cellular motion. Cells of wild-type and GldJ ΔC8 strains exhibited average locomotion velocities of about 2 μm/s and 0.4 μm/s, respectively (Fig. 2B, Supplementary video 3). The above results indicate that in the GldJ ΔC8 cells the conveyor belt may either be shortened or even depolymerized. In contrast, GldK R73S, M77L and N307S suppressor mutants of GldJ ΔC8 background did exhibit runs of reduced length, with the average locomotion velocities being about 0.89, 0.64, and 0.88 μm/s. Notably, the oscillatory back-and-forth motion was present along with the runs. The longest runs among the three evolved GldJ ΔC8 strains occurred within the N307S population (Fig. 2C, Supplementary video 4). In summary, the suppressor strains exhibited an improvement in swarming and single-cell motility as compared with the parent GldJ ΔC8 strain. This implies that point mutations in the T9SS rotor protein GldK, which has around 30% sequence similarity to GldJ (Fig. S1B), functionally restores, to a certain degree, the mostly non-operational conveyor belt of the parent GldJ ΔC8 strain.

To gather more information on the integrity of the conveyor belt, a mobile cell-surface adhesin, SprB, which is attached to the conveyor belt, was fluorescently labeled and its movement was tracked. Trajectory of a single SprB signal provided information about the structural integrity of the conveyor belt. As anticipated, wild-type cells exhibited long trajectories with SprB moving at about 2 µm/s, and they appeared to have fully formed, closed conveyor belts looping around cell-poles. In contrast, GldJ ΔC8 strain exhibited drastically short SprB trajectories characteristic of the oscillatory motion at about 0.5 µm/s, suggesting either immobilization or depolymerization of the conveyor belt, but N307S strain of GldJ ΔC8 background exhibited longer trajectories akin to the wild type. This indicates that the functionality of the GldJ conveyor belt is restored, at least, partially, because of the above-mentioned point mutations in the GldK ring of T9SS. In the latter case, the velocity of SprB was about 1 μm/s, half of that characteristic of wild-type cells, suggesting that the T9SS rotor to conveyor belt interaction was partially restored (Fig. 3, Supplementary video 5). To summarize, the detailed analysis of swarming motility and single-cell motility, facilitated by SprB trajectory tracking, provided evidence that the interaction between C-terminal region of GldJ and the three experimentally identified mutations of GldK is crucial for maintaining proper operation of the conveyor belt, which might enable the cell to search a larger area for nutrients. Additionally, the three GldK suppressor strains isolated from our screen also had a N178K mutation in a putative LuxR-like protein (*Fjoh_4220*) and a G223V mutation in an Acyl Carrier Protein (ACP) Synthase (*Fjoh_3810*). A detailed genetic, single-cell motility, and swarming analysis showed that *fjoh_4220* and *fjoh_3810* do not impact gliding motility (Fig. S2, Fig. S3, Supplementary text, and Supplementary videos 6-9).

**Figure 3.**
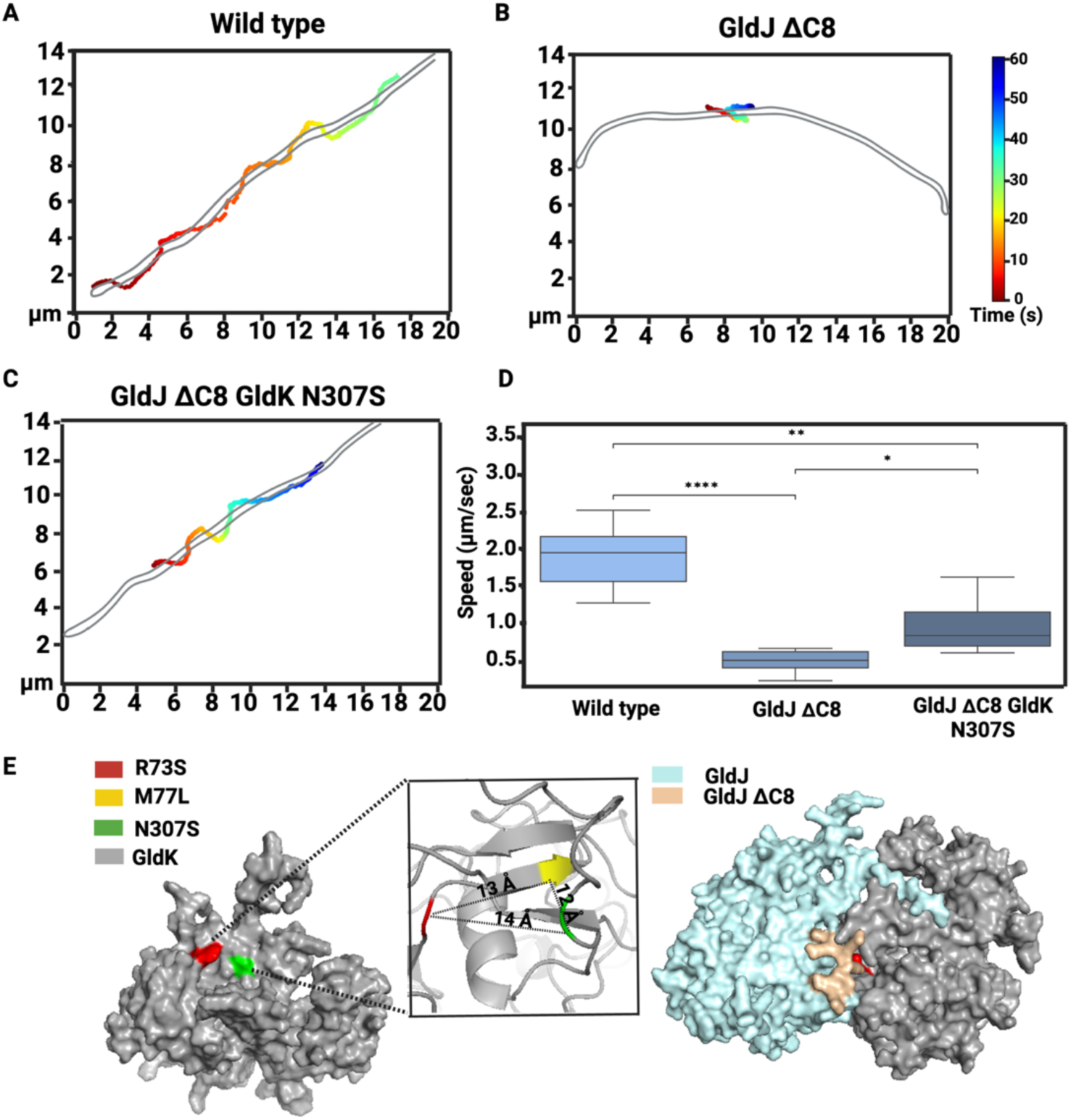
Motion of the cell-surface adhesin SprB is partially restored in GldK N307S carrying GldJ ΔC-terminal cells. **(A-C)** A representative trajectory of fluorescently labeled SprB moving along the conveyor belt of wild-type and mutant cells. **(D)** Box plot showing speed of at least six SprB molecules for each strain. Statistical significance was analyzed via student’s t test. P value: (*P < 0.05, 10^-3^ < **P > 10^-2^, ****P <= 10^-4^). **(E)** Predicted structure of GldK shows that the amino acids R73, M77, and N307 identified via the suppressor screen are in close vicinity. Docking of the conveyor belt protein GldJ (cyan) with the T9SS rotor protein GldK (gray) shows that the GldJ C8 region fits into a GldK cleft region that contains R73 and N307 with M77 in vicinity.

### Suppressor mutagenesis unlocked some of the CW-only T9SS motors

As evident by the data presented in Fig. 1, T9SS of the parent GldJ ΔC8 strain are “locked” in the CW-only rotational state. Because the evolved GldK point mutations partially restored cellular motility lost due to ΔC8 in GldJ (Fig. 2, Fig. 3), we tested if some of the T9SS motors in the evolved strains could rotate CCW. Indeed, the GldK point mutations influenced the rotational bias as well, with some T9SS motors of the evolved strains rotating CCW, albeit about 5-10 times slower than those having wild-type GldK. At such low rotational frequencies, it is tough to differentiate between rotational motion and Brownian fluctuations. Therefore, only the faster population of T9SS motors that exhibited at least one full revolution were further analyzed. The above criterion increased the statistical significance of our results, but, simultaneously, it reduced the overall population of T9SS motors that were subjected to the analysis. We found that in the R73S strain of GldJ ΔC8 background one out of six T9SS motors taken into consideration rotated CCW (Fig. S4). In the N307S strain the above was true for one out of sixteen T9SS motors (Supplementary video 10). Contrary to the wild-type units that exhibit CCW bias, T9SS of the evolved strains retained the CW bias of the parent strain. However, in both wild-type and experimentally evolved strains, cellular motility was observed irrespective of the direction of the rotational bias (Fig. 2). In both cases, a rotational bias (whether CCW or CW) of about 85 % to 95 % and about 5 % to 15 % noise contribution was associated with long runs that are characteristic of typical gliding trajectories. In the absence of noise, *i.e*, 100 % CW bias characteristic of the GldJ ΔC8 strain, the cellular motility is diminished (Fig. 1B, 2C).

In the ideal case, a gliding cell would be equipped with about 8 to 12 T9SS units rotating in unison^34,26^. If one of the units were to rotate in the opposite direction, it would contribute about 8 % to 12.5 % to the noise, which is in agreement with the CCW:CW bias observed in our tethered-cell data (Fig. 1B). Given that multiple T9SS motors control a single conveyor belt, a random change in the rotational direction of any motor, due to inherent noise, could lead to slipping of the conveyor belt. This slippage might then cause disorientation and potentially change the gliding direction of the cell. Within seconds, the slipped conveyor belt might diffuse and re-engage with T9SS motors rotating concertedly, leading to restoration of smooth gliding behavior, albeit in a new direction. The observation that gliding cells follow straight trajectories interspersed with sudden turning events (Fig. 2C) further supports the mechanism suggested above. In their natural environments, gliding cells release enzymes to decompose complex polysaccharides such as chitin and cellulose found in insoluble food sources^35,36^. Their long gliding movements, interspersed with sudden turns, may enhance their ability to locate these food sources. This behavior suggests evolutionary pressure for a strong CCW rotational bias, with significant CW noise. In the lab, we have observed this pattern in wild-type strains (Fig. 1B, 2C) and its reversal in evolved strains, where a prominent CW bias is coupled with noticeable CCW noise (Fig. S4). The observed “inverse state,” characterized by slower cell movement, suggests that the interaction energy between T9SS units and the conveyor belt differs between wild-type and evolved strains.

A predicted AlphaFold tertiary structure of GldK shows that the residues R73, M77, and N307 are packed close together in an arrangement which resembles a cleft, with the R73 to N307, R73 to M77, and M77 to N307 inter-residue distances being 14 Å, 12 Å, and 13 Å, respectively (Fig. 3E). Furthermore, GldK has been shown to facilitate the oligomerization and complex formation of GldJ^23^. Consequently, Molecular docking of GldJ and GldK was performed using HDOCK^37^. One of the models, which has an 86% accuracy score, indicates that the C-terminal region of GldJ fits into the cleft formed by GldK and aligns with the genetic data (Fig. 3E). Therefore, it was selected for further analysis. MD simulations performed on the docked GldJ-GldK model revealed that deleting the C8 region of GldJ causes the cleft region of GldK, which contains residues R73, M77, and N307, to become more dynamic and expand towards the area previously occupied by C8 of GldJ (Fig. S5). This explains why mutations in the ring protein GldK can compensate for the loss of the C-terminal region of the conveyor belt protein GldJ. This provides further evidence that the phenotypes described above might be due to changes in the interaction energy between the T9SS ring and the conveyor belt. Interestingly, the combination of GldK mutations R73S, M77L, and N307S in a GldJ ΔC8 background does not restore either swarming on agar or single-cell motility over a glass surface (Fig. S6 and supplementary text). This suggests that these three GldK point mutations together might completely hinder the interaction of the rotor with the conveyor belt. Any one of the three GldK mutations is sufficient to alter the function of the cleft region of GldK and consequently its interaction energy with GldJ ΔC8.

### Molecular simulations suggest that GldLM stator units rotate in the clockwise direction

In a gliding cell, the GldKN/PorKN ring interacts with the GldLM/PorLM stator units ^38^, which harness pmf to power the rotation of the T9SS^27^. Recent research has demonstrated that in flagellated bacteria, the MotAB components of the bacterial flagellar motor rotate CW, driving the rotation of C-rings of the bacterial flagellar motor^39^. The proton-conducting GldLM stator units have an asymmetric 5:2 structure similar to MotAB. However, despite this conceptual similarity, the actual rotational dynamics of GldLM stator units at the atomistic level have not yet been explored. Given that MotAB components rotate unidirectionally CW^39^, analyzing whether GldLM stator units are bidirectional or unidirectional may help elucidate if GldLM play any role directional switching of T9SS. To investigate this, we employed MD simulations to model the rotational dynamics of GldLM.

The probability of CW vs CCW GldLM rotation was estimated via MD. Notably, GldLM has a characteristic structural asymmetry with two GldM chains distributed among five GldL subunits. The latter feature precludes uniformity of GldL-GldM interactions, and it has been suggested that a single GldM chain can only be engaged into a channel-specific ion pairing involving protonated ARG 9 of GldM and deprotonated GLU 49 of GldL at a time^25^. This ion pair is implicated in transporting protons and generating a proton motive force. With the PDB all-atom macromolecular structure featuring GldLM symmetry mismatch as an input (see PDB: 6YS8), three different MD simulations were performed on the GldLM stator units.

We began with an all-atom equilibrium MD simulation where the non-bonded interaction energies were monitored across the simulation trajectory to explore the contacts between ARG 9 of GldM and GLU 49 of GldL, where GldL is either in CCW or CW orientation. We obtained the mean values of -57.5 and -94.6 kcal/mol for the CCW-orient GLU 49 and CW-oriented GLU 49 on GldL, respectively (Fig. 5B). The rotation of transmembrane helices of GldM within GldL is inaccessible using traditional MD techniques, hence 20 replicas of 10 ns-long steered MD (SMD) simulations were explored. The results are indicative of a significant relationship between average work and rotational directionality preference, suggesting a bias towards CW (Fig. 5C).

Prompted by the apparent differences in work profiles, we sought to obtain a more reliable description of the CCW vs. CW occupancies in GldM through Modeling Employing Limited Data or MELD-guided MD simulations^40^. We began with a configuration where the ARG 9 residue was equidistant from the CCW and CW deprotonated GLU 49 of GldL to avoid any rotational bias in our model. Upon convergence, our MELD simulation revealed that around 82% of ARG tend to be spontaneously located in the immediate vicinity of the CW-oriented GLU, in the absence of any directive steering force throughout the simulation. Thus, as opposed to the (CCW biased) GldK ring of T9SS, the proton conducting GldM-GldL transmembrane helix complex of T9SS have a CW bias (Fig. 5D). In our MD simulations, GldLM ion channels rotated mostly CW, and we observed no transitioning to CCW rotation. Consequently, the MD simulations revealed that the directional bias change in the T9SS cannot be ascribed to a change in rotational directionality of the GldLM ion channels that drive this system.

## DISCUSSION

Genetic data, cell tethering assays, and MD simulations combined with biochemical and structural evidence for the interactions between T9SS and motility proteins such as the GldKN ring and GldLM stator units and between GldJ and GldK from various organisms^23,25,27,34,41,42^, suggest that the T9SS GldKN ring, the GldJconveyor belt, and the GldLM stator units function as three interlocking mechanical gears. We term this interaction a tri-component gearset. In this system, the GldLM stator unit serves as the primary gear and driver, utilizing proton motive force. The stator unit drives the rotation of the middle gear, the GldK ring, which in turn propels the outer component, the GldJ conveyor belt. Cryo-ET has revealed that the GldJ conveyor belt forms a closed loop, with multiple such loops observed within a single cell^23^. When examined in the 2D plane, the outer loops of the tri-component gearset appear as linear conveyor belts (Fig. 5a). Our findings suggest that the GldJ conveyor belt may influence the rotational directionality of the GldK-containing T9SS ring, while the GldLM stator units, responsible for generating force, rotate CW unidirectionally, propelling the middle gear (GldK ring) of the tri-component gearset.

Here, we advanced a working model by estimating the interaction energy between GldJ and GldK monomers using MD simulations. The simulation trajectories were analyzed to assess van der Waals and electrostatic interactions between GldJ and GldK at every timestep over a 100 ns simulation period. We observed that, for the monomers, the total non-bonded GldJ-GldK interaction energy decreased by about 25% upon removal of the 8 C-terminal amino acids of GldJ (Fig. S7). We further observed a significant conformational change in the tertiary structure of monomeric GldK upon interaction with GldJΔC8 compared to wild-type GldJ (Fig. 5B and Fig. 5C). Additionally, RMSD values for GldK reformations increased by 10 Å after the interaction interface with GldJ was altered by the C8 deletion (Fig. S8). In the docked model, the distance between the centers of mass of GldK and both forms of GldJ increased by 7.5 Å during the MD simulation (Fig. S9), reinforcing our inference that the tertiary structure of GldK undergoes conformational changes after the deletion of the C8 region of GldJ. In stark contrast, the structural changes in GldJ lacking the eight C-terminal amino acids are similar to those in wild-type GldJ (Fig. S10), thus providing an internal control for the MD simulations. An overlay of GldK monomer from the GldJΔC8 model with the wild-type GldK monomer shows several regions in which the conformation of GldK changes > 1 nm (Fig. 5D). Interestingly, recent cryo-ET studies revealed that the T9SS GldKN (PorKN) ring of *P. gingivalis* adopts three distinct conformational classes, forming angles of 90°, 70°, and 50° with the outer membrane^27^ (Fig. 5F). This supports the results of our MD simulations, which also suggest that GldK can exist in multiple conformational states. In future, additional insights into the multimeric structure of the gliding machinery will a detailed understanding of how the conveyor belt regulates T9SS directionality.

Building on structural insights from *P. gingivalis* and extending our simulations to a larger scale, we propose a model where the interaction energy between GldJ and GldK may affect the size of the GldK ring, prompting a conformational shift once the C8 segment of GldJ is removed. We suggest that shifts in rotational directionality in T9SS might occur due to this conformational change of GldK. Tethered cell assays indicate that the wild-type T9SS predominantly rotates CCW. Therefore, the CW rotation of the GldLM stator units in wild-type cells likely propels near the outer edge of the GldK ring. This setup is comparable to a mechanical system where a CW rotating driving gear moves a second gear. In this case, the second gear (GldK ring) rotates CCW which is opposite of the CW rotation of the GldLM driving gear (Fig. 5G). According to theux model, the rotation of the GldK ring, either CW or CCW, depends on where the CW rotating GldLM stator units interact with the GldK ring in a manner reminiscent to that occurring between the MotAB stators and the flagellar C-ring^39^. At a cursory glance, the suggested mechanism behind directional switching of T9SS presented here seems similar to how the rotation direction of the bacterial flagellar motor is regulated^43^. However, the proteins composing the T9SS, and gliding machinery bear no sequence resemblance to those forming the bacterial flagellar motor, indicating an instance of convergent evolution. The structural resemblance between the transmembrane domains of GldLM and MotAB, combined with our MD simulations of GldLM (Fig. 4) implies that the underlying rotary mechanism might operate similarly in these two different types of motors^25^. Unlike the bacterial flagellar motor, which is regulated by soluble cytoplasmic signaling proteins, the outer-membrane-associated conveyor belt protein GldJ appears to uniquely influence the conformation of the T9SS ring and, consequently, the rotational directionality of T9SS. If our model is correct, it suggests that despite significant differences in mechanical structure, protein sequences, and biological functions between the T9SS and the bacterial flagellar motor, their regulation appears to adhere to similar principles, indicative of convergent evolution. In both instances, changes in ring conformation - resulting from structural adjustments within the flagellar FliG ring^43^ or from conveyor belt-driven conformational remodeling of the T9SS GldK ring, affect the positioning of the MotAB or GldLM stator units, respectively. This shift in positioning, in turn, causes the direction of rotation of the FliG or GldK ring to reverse. Structural studies of T9SS in *P. gingivalis* show that GldK (PorK) arranges into a ring^44^, and the core T9SS proteins forming the ring and stator units in both *P. gingivalis* and *F. johnsoniae* share a high degree of homology^25^. Our model consolidates current knowledge and serves as a springboard for future endeavors. Alternatively, it is also possible that the bend of GldM could be altered by a reduction of interaction energy between GldJ and GldK – which might lead to a change in its interaction site with the GldK ring as observed by cryo-ET^27^. By elucidating the multimeric structure of the gliding machinery, capturing higher-resolution views of the polymeric conveyor belt, and performing in-depth biophysical analyses of T9SS rotation of various mutants, we anticipate the emergence of more robust models in the future.

**Figure 4.**
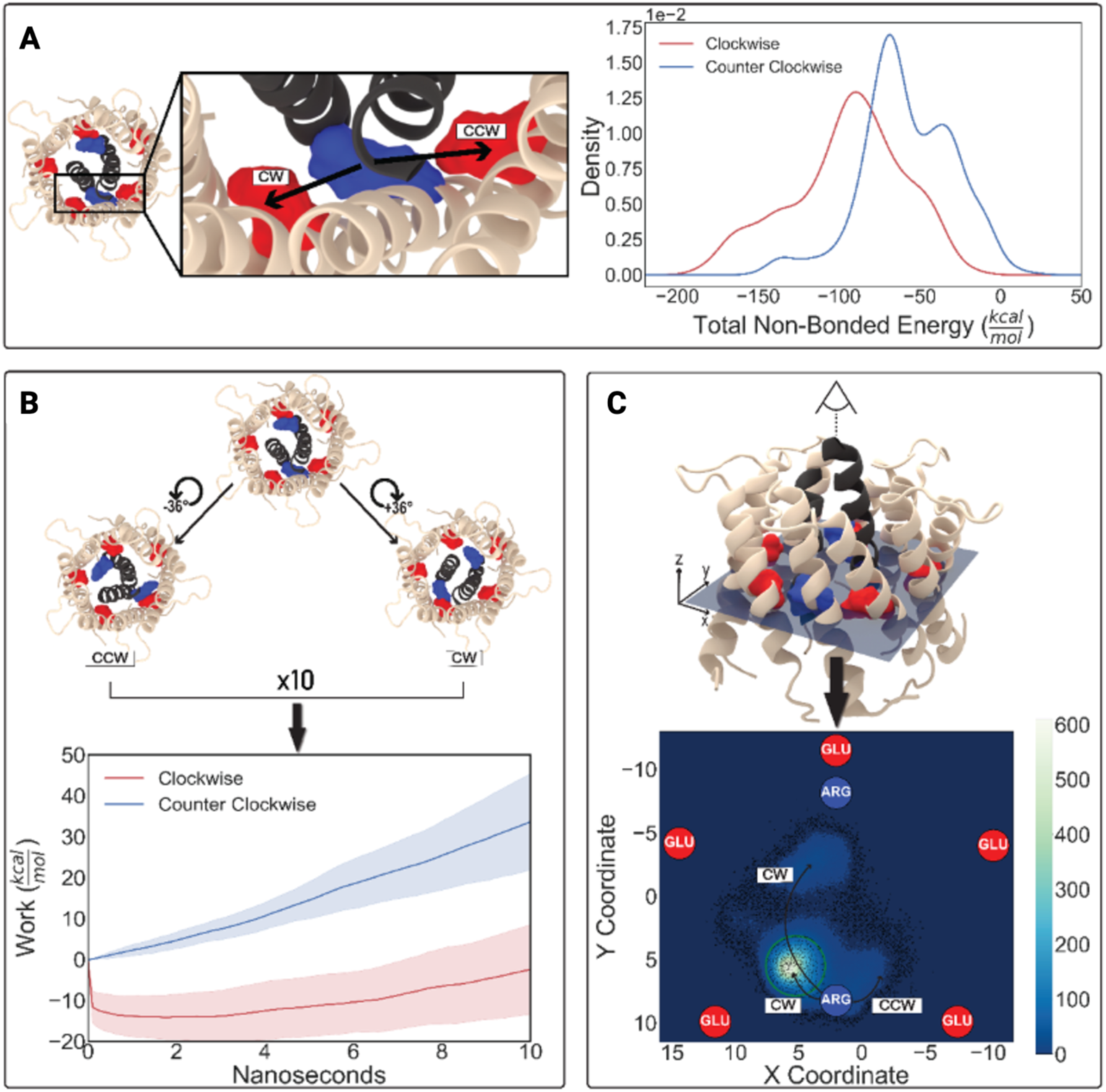
The driver gear, GldM, in the tri-component gearset rotates unidirectionally in a CW direction. **(A)** Equilibrium MD simulations and non-bonded energy calculations show that the central arginine residue of GldM interacts more favorably with the glutamate residue of GldL in the CW direction. The Kernel Density Estimation (KDE) method was used to estimate the probability density function of the non-bonded energy from the simulation trajectory. **(B)** Central: Unaltered equilibrium configuration of the initial structure, excluding the membrane for the sake of clarity. Left: Structure after a CCW GldM trans-membrane helix (TMH) rotation of -36°. Right: Result after +36° CW rotation. The plot illustrates the non-equilibrium energetics from ten consecutive SMD simulations (360° total in each direction), which reveal a lower work value for CW GldM TMH rotation. **(C)** Three-dimensional representation of the central structure shown in panel C. A top-down view highlights the plane used for the statistical analysis of post-MELD simulations. The spatial coordinates of the unbound arginine residue are mapped in a contour plot, with lighter shades indicating higher density, which suggests an unbiased preference for CW rotation. The arrows indicate the trajectory of the positional displacements of the unbound arginine. A green circle superimposed on the plot serves as the boundary where the probability of a datum falling within the enclosed region was computed.

**Figure 5.**
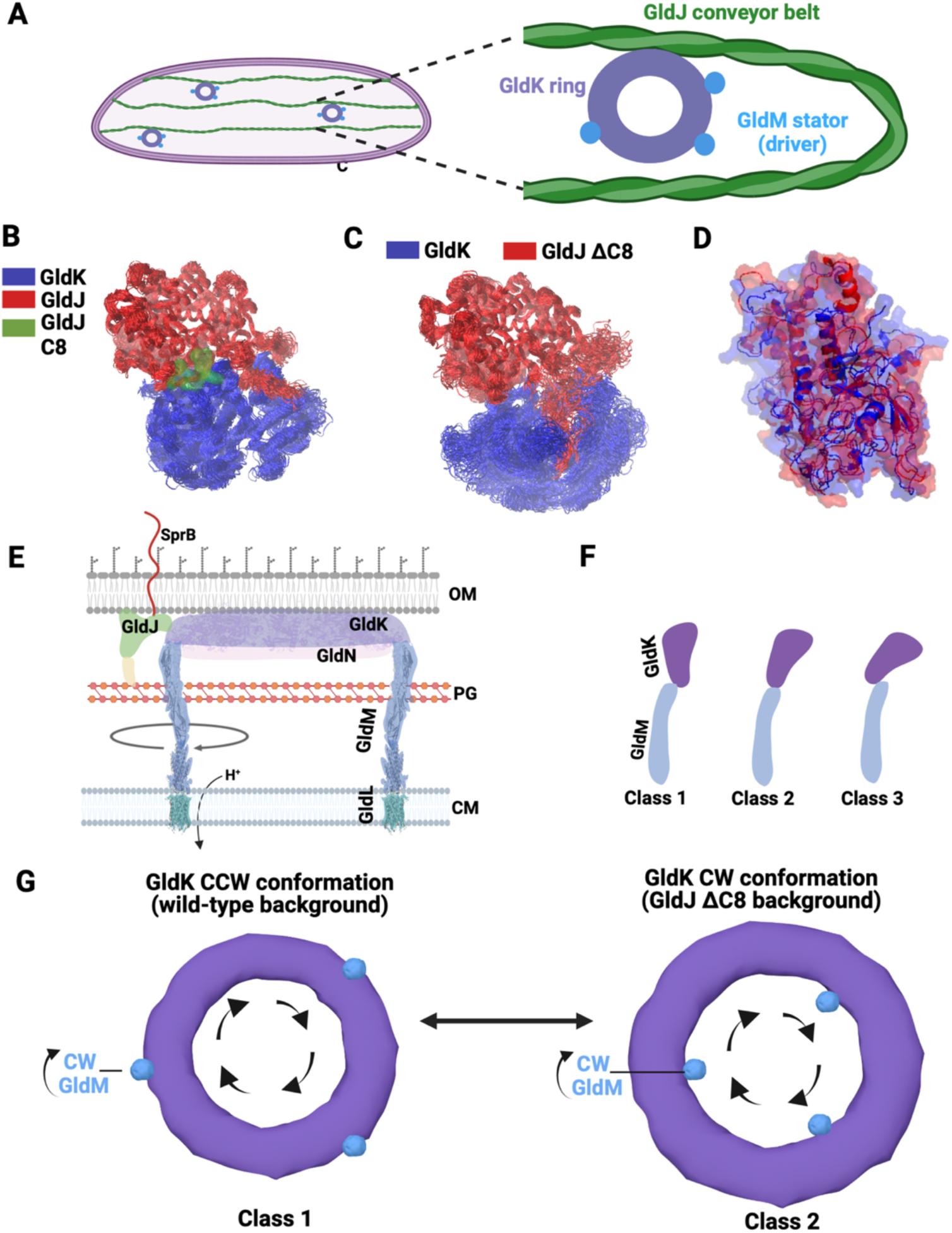
A model for directional switching of T9SS. **(A)** A cartoon showing the tri-component gearset with GldM stator unit as the driver of the T9SS ring and conveyor belt. **(B)** An overlay of frames collected for the final 50 ns of the MD simulation for the wild-type GldJ (red) and GldK (blue) system with the GldJ C8 region highlighted in green. **(C)** Upon deletion of the C8 region of GldJ, an appreciable expansion of GldK is observed in contrast to the wild-type system shown in panel B. **(D)** An overlay of the GldK from the wild-type and GldJΔC8 systems depicts conformational changes that are > 1nm. **(D)** A cross-sectional cartoon of the gliding machinery shows the proton motive force (pmf) powered driver gear, GldM, rotating unidirectionally clockwise (CW) and pushing the T9SS ring. This action results in the movement of the GldJ conveyor belt and SprB adhesin either towards the viewer or away from the current image plane. Images are not to scale. **(E)** A cartoon summarizing the cross-section of different conformational states (class1, 2, and 3) of T9SS ring as observed by cryo-ET^27^. **(F)** A top down view of class 1 and class 2 T9SS motors as observed by cryo-ET^27^ supports a physical model of T9SS directional switching. When the wild-type conveyor belt protein GldJ interacts strongly with the GldK ring T9SS maintains a conformation where GldM pushes the outer periphery of the GldK ring, causing T9SS to rotate CCW. Conversely, when the interaction energy between GldJ and GldK is reduced (GldJΔC8), the GldK ring changes conformation. Consequently, GldM pushes near the inner periphery of the GldK ring, resulting in CW rotation of the T9SS.

Overall, our findings suggest an additional role for the GldJ conveyor belt, particularly of its C-terminal region, which may function as a molecular switch. Powered by proton motive force, the T9SS stator units play a role akin to that of the driving motor in a mechanical conveyor system. The GldK ring, on the other hand, is akin to the gear that translates rotational energy of the motor into linear motion of the conveyor belt. Meanwhile, the GldJ loop represents the conveyor belt itself, but with an added layer of complexity - unlike a passive mechanical conveyer belt that we commonly encounter in our daily life, GldJ plays a dual role; in addition to be passively driven by forces exerted on it by the GldK gear, it also feeds back into the system to influence and control the rotational directionality of its driving gear itself. In traditional human-engineered mechanical conveyor systems, a series of motors and gears propel the belt along a predetermined path. However, more advanced systems, known as flexible or cognitive mechanical conveyors, are equipped with routing algorithms that adjust the direction of motion of the conveyor belt to adapt to varying requirements^45^. Shaped by evolutionary pressures, the bacterial gliding machinery described here represents a unique example of a conveyor belt equipped with sensory feedback. The data presented here demonstrate that bacteria have evolved a smart conveyor belt system capable of reciprocally altering the rotational bias of its driver, thereby potentially functioning as a dynamic and adaptive sensor or controller. Our model establishes a foundation for future investigations into the molecular mechanisms of this conveyor belt system and its adaptations to shifting environmental conditions. In doing so, this study paves the way for unraveling the intricacies of bi-directional feedback controllers in bacterial cells, whose inner workings resemble a controllable biological snowmobile.

## METHODS

### Strains, media, and culture conditions

For standard culture maintenance, genetics, and tethered cell analysis, wild-type *F. johnsoniae* CJ 1827^46^ cells were grown in Casitone Yeast Extract (CYE) broth (10 g casitone per liter, 5 g yeast extract per liter, 1 M Tris-HCl pH 7.5) at 30°C with 1.5% agar. *Escherichia coli* cells used for preparation of genetic constructs were grown in Luria Bertani medium (10g Tryptone per liter, 10g Nacl per liter, 5g yeast extract) at 37°C. Unless otherwise noted, antibiotics were used at the following concentrations: ampicillin (100 μg/ml), erythromycin (100 μg/ml), streptomycin (100 μg/ml), and cephalexin (75 μM or 26 μg/ml).

### Measurement of swarming motility on agar

*F. johnsoniae* cells were grown in PY2 broth (2 g peptone per liter, 0.5 g yeast extract per liter, pH 7.3) overnight at 25°C with shaking at 50 rpm. Cells were then washed once with PY2 medium and adjusted to an OD_600_ of 0.1. A 2 μl sample was spotted onto the center of a 1% PY2 agar petri plate, which was then incubated at 25°C for 48 hours. After incubation, the plates were imaged using an Amscope LED Trinocular Zoom Stereo microscope (SM-2TZZ-LED-18M3) with a camera (AmScope 18MP Color CMOS, C-Mount Microscope Camera, MU1803).

### Analysis of gliding motility on a glass surface

*F. johnsoniae* cells were grown overnight in CYE broth. Then, 50 μl of cells were inoculated in 5 ml of MM and grown for 6 hours at 25°C with shaking at 50 rpm until the OD_600_ reached 0.4. After that, 45 μl of the culture was introduced into a tunnel slide, incubated for 5 minutes, washed with 40 μl of MM, and imaged with a Nikon Optiphot phase contrast microscope fitted with Thor lab CMOS camera (DCC1545M). Videos were recorded at 15 fps. Custom Python scripts were used to analyze the trajectory and speed of the cells.

### Suppressor mutagenesis

*F. johnsoniae* strains lacking 8 amino acids from C-terminal of GldJ (GldJ553/CJ2443) were grown in MM at 30°C for 24 hr. A volume of 20 μl was taken and spotted at the center of twenty different PY2 plates, dried for 20 min, and kept for incubation (lid up) in a 100% relative humidity chamber at 25°C for 3-4 days. Spreading colonies flared out on PY2 plates (Fig. S1). These flares were streaked for isolation and single spreading colonies were selected. Genomic DNA was isolated from the cell using GeneJET Genomic DNA Purification Kit (K0722) and the whole genome was sequenced (Supplementary data S1). As a result, three individual strains, which had the following mutations - GldK R73S (strain name: FJASU1), GldK N307S (strain name: FJASU3), GldK M77L (strain name: FJASU4) flared out as suppressor mutants from (GldJΔC8/CJ2443). All spreading colonies also had the G223V mutation in ACPsynthase (*fjoh_3810*) and N178K mutation in putative LuxR (*fjoh_4220*). Separate mutagenesis of each gene revealed that *fjoh_3810* and *fjoh_4220* did not influence this phenotype (Fig. S2, S3).

### Construction of point mutations

To obtain the 2 kb fragment of *gldK* from the suppressor mutants FJASU1, FJASU3, and FJASU4, Phusion high-fidelity PCR master mix (Thermo Scientific) was used with primers PASU52 and PASU53. The resulting fragment was then digested with BamHI and SalI restriction enzymes and cloned into the pRR51 plasmid previously digested with the same enzymes. This plasmid was then introduced into *F. johnsoniae* through triparental conjugation and mutants were isolated as described previously^46^. Briefly, a mixture of PRR51 containing *E. coli* strain, helper *E. coli* strain, and CJ1827 was spotted on a CYE agar plate containing CaCl_2_. Next day, the mixture was plated with dilutions on a CYE plate containing erythromycin to select for cells that had undergone the first recombination step. The erythromycin resistant colonies were cultured overnight in CYE broth and then plated with dilutions on a PY2 agar plate containing streptomycin to select for cells that had undergone the second recombination step. Point mutations in ACP and LuxR were generated via a similar method with the following exceptions – primers PASU26 and PASU27 were used to amplify ACP synthase region while primers PASU50 and PASU51 were used to amplify putative LuxR region. All primers and plasmids used in this study are in Tables S1 and S2. All mutations were confirmed by Sanger sequencing (Azenta life sciences) All sequencing data is available on dropbox (link).

### Immunofluorescence labelling of SprB and imaging via TIRF microscopy

Single colonies of the *F. johnsoniae* GldK N307S in GldJ ΔC8 strain (FJASU19) were grown overnight in MM at 25°C with shaking at 50 RPM. From the overnight saturated culture, 100 µL was used as an initial inoculum to subculture 5 mL of MM containing cephalexin and incubated at 25°C with shaking at 50 RPM, resulting in the generation of elongated cells. After 6 hours, 100 microliter of culture was washed by centrifugation at 5000 g for 1 min. The resulting pellet was resuspended in 45 μl of MM and 5 microliters of 1:10 diluted purified anti-sprB antibody was added. Anti-SprB antibody^47^ was purified using Melon Gel IgG Spin Purification Kit (Thermo Scientific, 45206) as described previously^1^. After incubation at room temperature for 10 min, cells were washed again via centrifugation at 5000 g for 1 min and resuspend in 45 μl MM. The cells were treated with 2 µL of secondary antibody, Alexa Fluor 555-labeled anti-rabbit IgG (Abcam, Cambridge,UK), and incubated at 25°C for 10 minutes. The cells were washed thrice at 5000 g and were pipetted into a tunnel slide. Cells were incubated for 5 min at RT and washed gently with MM in the tunnel slide. Imaging was performed on ONI Nanoimager microscope (Oxford Nanoimaging) equipped with 405 nm diode pump solid state lasers. Optical magnification was provided by a Å∼100 oil-immersion objective (Olympus, 1.4 numerical aperture) and images were acquired using an ORCA-Flash4.0 V3 CMOS camera (Hamamatsu). TIRF images were acquired using a 20-ms exposure time, with the 561-nm photoactivation laser at 2% power.

### Tethered cell assay

The cells were grown overnight in MM at 25°C with shaking at 50 RPM. From the overnight culture, 100 µL was sub-cultured into fresh 5 mL of MM media and incubated at 25°C with shaking at 50 RPM. When the culture reached an OD_600_ of 0.4, cell tethering was performed as previously described^3^. Briefly, a polyethylene tubing with an inner diameter of 0.58 mm was used to shear cells by passing 500 µL of the culture 50 times through the tubing using 1 mL syringes with 23 G stub adapters. (Instech PE-50 tubing, Instech Luer stub LS23S, Fisherbrand sterile plastic syringe). The sheared cells were washed with MM by centrifugation at 5000 g for 1 min. A volume of 40 µL of this sample was incubated with anti-sprB antibody diluted (1:10) for 20 minutes at room temperature. After another wash at 5000 g for 1 minute in MM, the cells were pipetted into the tunnel slide. After 5 minutes, 200 microliters of MM were gently flowed through the tunnel slides multiple times until residual antibody was washed away and freely rotating cells were observed.

### Preparation of the MD model

The initial model of the membrane protein system was prepared using CHARMM-GUI membrane builder. The protein structure, consisting of five GldL subunits and two GldM transmembrane helices^25^, was uploaded in PDB format, and the system was solvated in a preequilibrated phosphatidylcholine (POPC) lipid bilayer. TIP3P water molecules were added to the system, along with 0.15 M NaCl, to reproduce cellular conditions.

### Equilibrium MD

All-atom molecular dynamics simulations were performed using Nanoscale Molecular Dynamics (NAMD) 3.0 using the CHARMM36m force field for protein, lipids, and ions. Periodic boundary conditions were implemented in all dimensions. The system was energy-minimized followed by an equilibration phase under NPT conditions at 300 K and 1 atm for 1µs.

### Steered MD

Non-equilibrium molecular dynamics simulations were carried out using a spin angle CV; one of a two-part decomposition of a full rotation CV effectively measuring the angle of rotation of the GldM TMH around the z-axis. A total of 20 simulations were executed, evenly bifurcated into two directional categories. The rotation parameter was established at ± 36°. Negative magnitudes denote a counterclockwise torsional movement, whereas positive magnitudes signify a clockwise torsion. Subsequently, after ten iterations, the GldM TMH underwent a complete rotation of ±360°. All simulations were performed using NAMD3 software using the CHARMM36m force field for protein, lipids, and ions. Periodic boundary conditions were implemented in all dimensions. The system was energy-minimized followed by a non-equilibration phase under NPT conditions at 300 K and 1 atm. Each simulation was run for 10 ns.

### Modeling Employing Limited Data (MELD)

MELD is a physics-based, Bayesian computational technique leveraging Open Molecular Mechanics (OpenMM) MD software. The simulation schematic involved using a 16-replica parallelization assigning one GPU per replica. The individual simulations spanned a duration of 250 nanoseconds each, with exchange attempts initiated at regular intervals of 10 picoseconds respectively. There was an implemented linear temperature range from 300 to 380 K across the replica ladder i.e., 300 K at replica 1, 380 K at replica 16 and uniform temperature interpolation for the remaining replicas. To effectively model solvation effects in our system, we utilized the Generalized Born implicit solvent model (GB-Neck2).

We chose the Boltzmann distribution as the prior, as it provides an unbiased view of the system’s conformational landscape. The likelihood distribution includes all external information, manifested as flat-bottom harmonic potentials, and comprised of non-negative constraints on geometric degrees of freedom. The external information was incorporated from generated contact maps between selected residues whose specificities are outlined below: We applied distance constraints to the heavy atoms within GldL with an absolute confidence of 100%, ensuring the inclusion of every computed contact; we imposed identical constraints within the GldM TMH. These heavy atom constraints maintained the structural integrity of the model especially at high-temperature replicas. There were no restraints imposed between GldM and GldL therefore allowing flexibility and rotation. The prior and likelihood constituted the posterior distribution which was ultimately sampled during MELD.

## Acknowledgements.

A. Shrivastava was supported by an NIH-NIGMS MIRA award R35GM147131. A. Singharoy was supported by an NSF CAREER award MCB-1942763. The MD simulation used an Oak Ridge Leadership Computing Facility, supported an award from the Office of Science, Department of Energy DE-AC05-00OR22725. We thank Mohammed Kaplan, Grant J. Jensen, and Stefano Maggi for helpful comments on the manuscript. We thank Pushkar P. Lele for helpful suggestions and Rizal F. Hariadi for help with TIRF microscopy. The images were created using biorender.com.

## Author contributions

AT generated the mutants and performed the cell tethering and TIRF assays. AT and ECH collected and analyzed the cell-motility data. A. Shrivastava generated the initial suppressor strains. JAM and A. Singharoy performed the MD simulations. A. Shrivastava and AT wrote the manuscript.

## Data Availability

Custom Python codes for image analysis and example data sets are freely available on our GitHub. The molecular dynamics models, and additional images are available upon request. All other study data are included in the article and/or supporting information.

## SUPPLEMENTARY FIGURES

**Figure S1.**
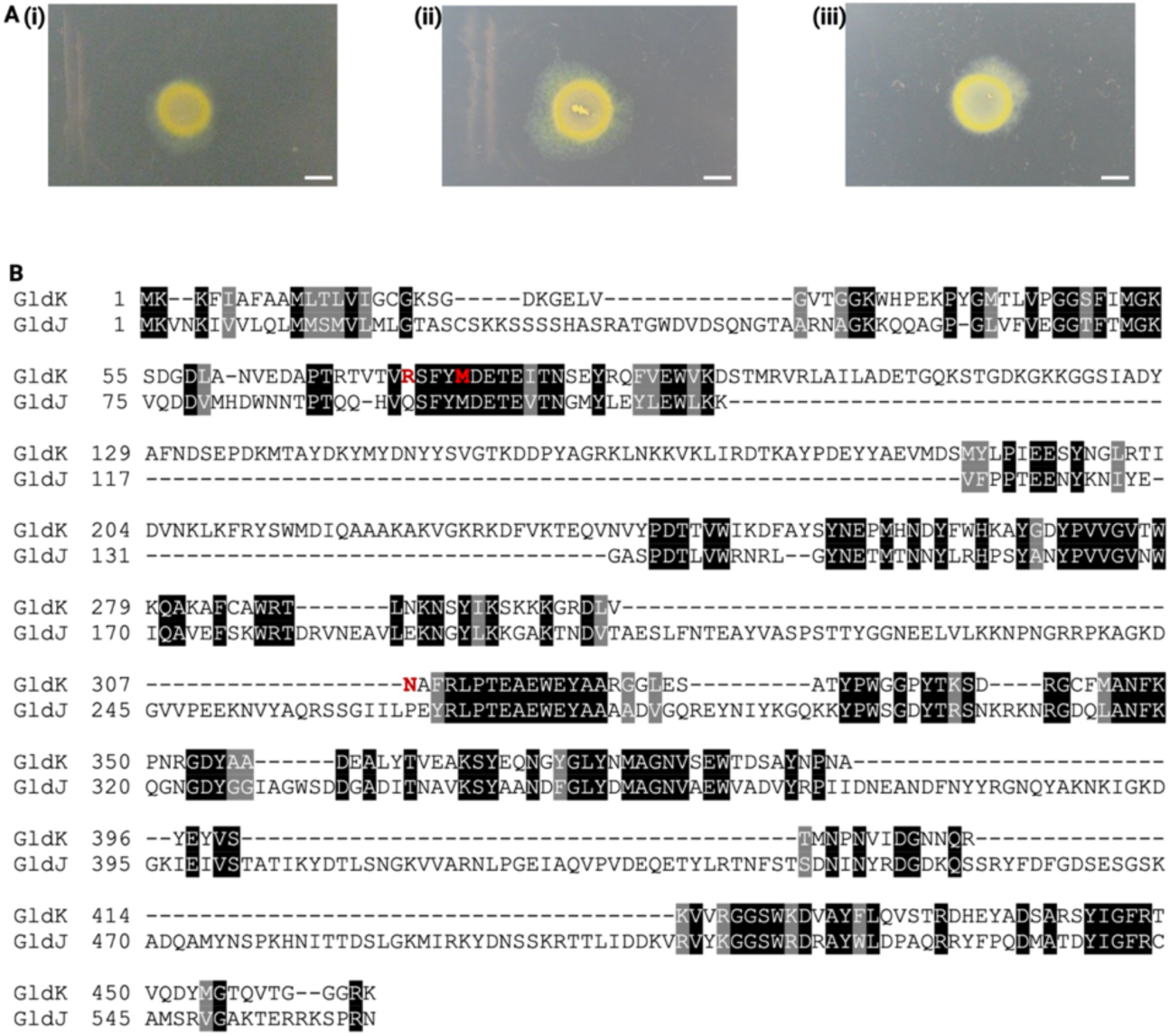
(A) Images from the experimental evolution screen of GldJ ΔC8 which lead to the following suppressors that had the ability to swarm on agar (i) GldK R73S (FJASU1) (ii) GldK N307S (FJASU3) and (iii) GldK M77L (FJASU4). Scale bar, 3 mm. (B) Multiple sequence alignments of *F. johnsoniae* GldJ and Gld

**Figure S2.**
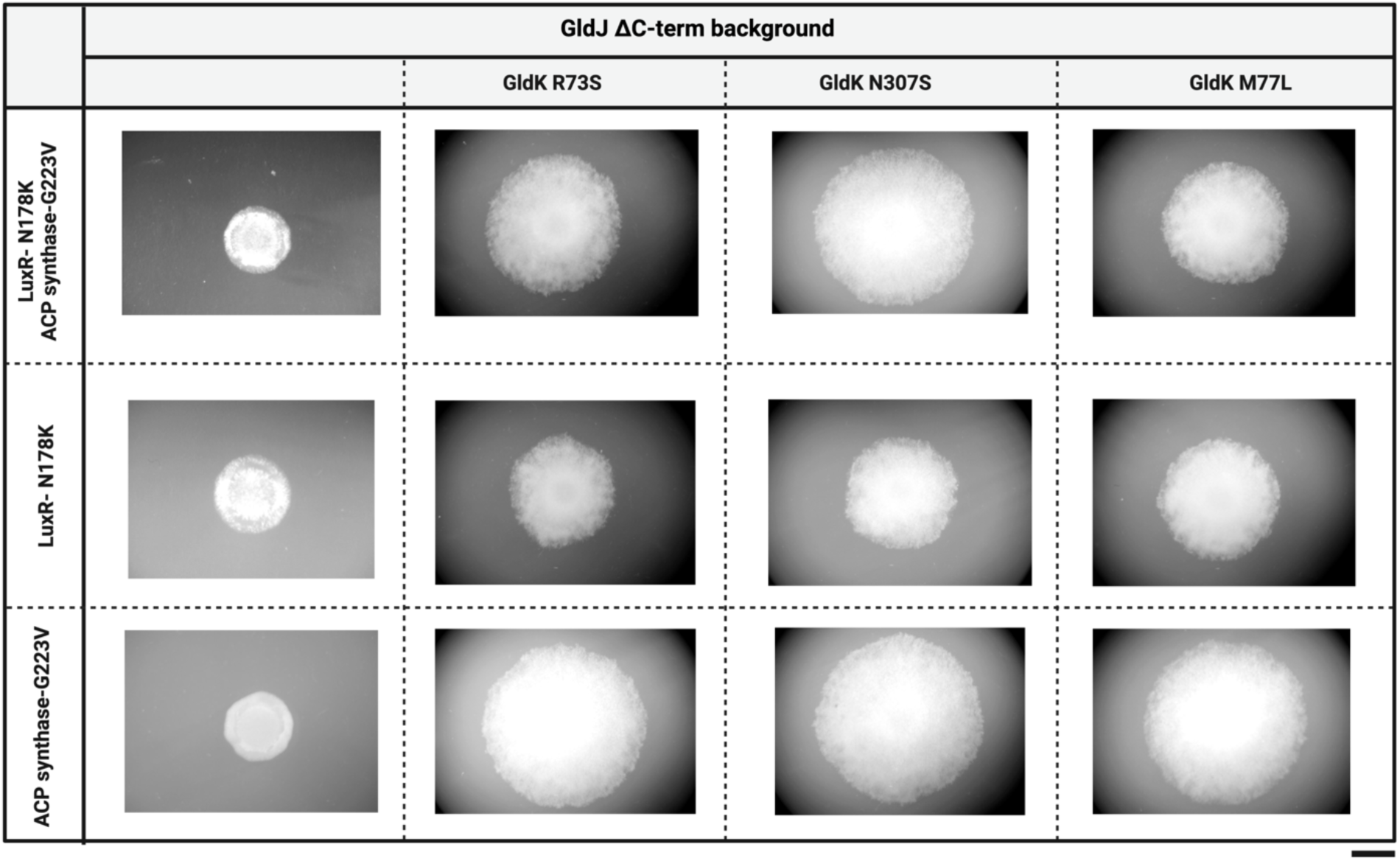
The evolved strains containing either GldK R73S, N307S or M77L restore the swarming motility of the parent GldJ ΔC8 (GldJ ΔC-term background) on PY2 agar plates. Scale bar, 6 mm. Whole genome sequencing revealed that evolved strains also had secondary point mutations in genes encoding an ACP synthase (G223V) and LuxR-like protein (N178K). Individual and combinatorial mutagenesis of ACP synthase and LuxR showed that these two genes do not play a role in restoration of swarming motility.

**Figure S3.**
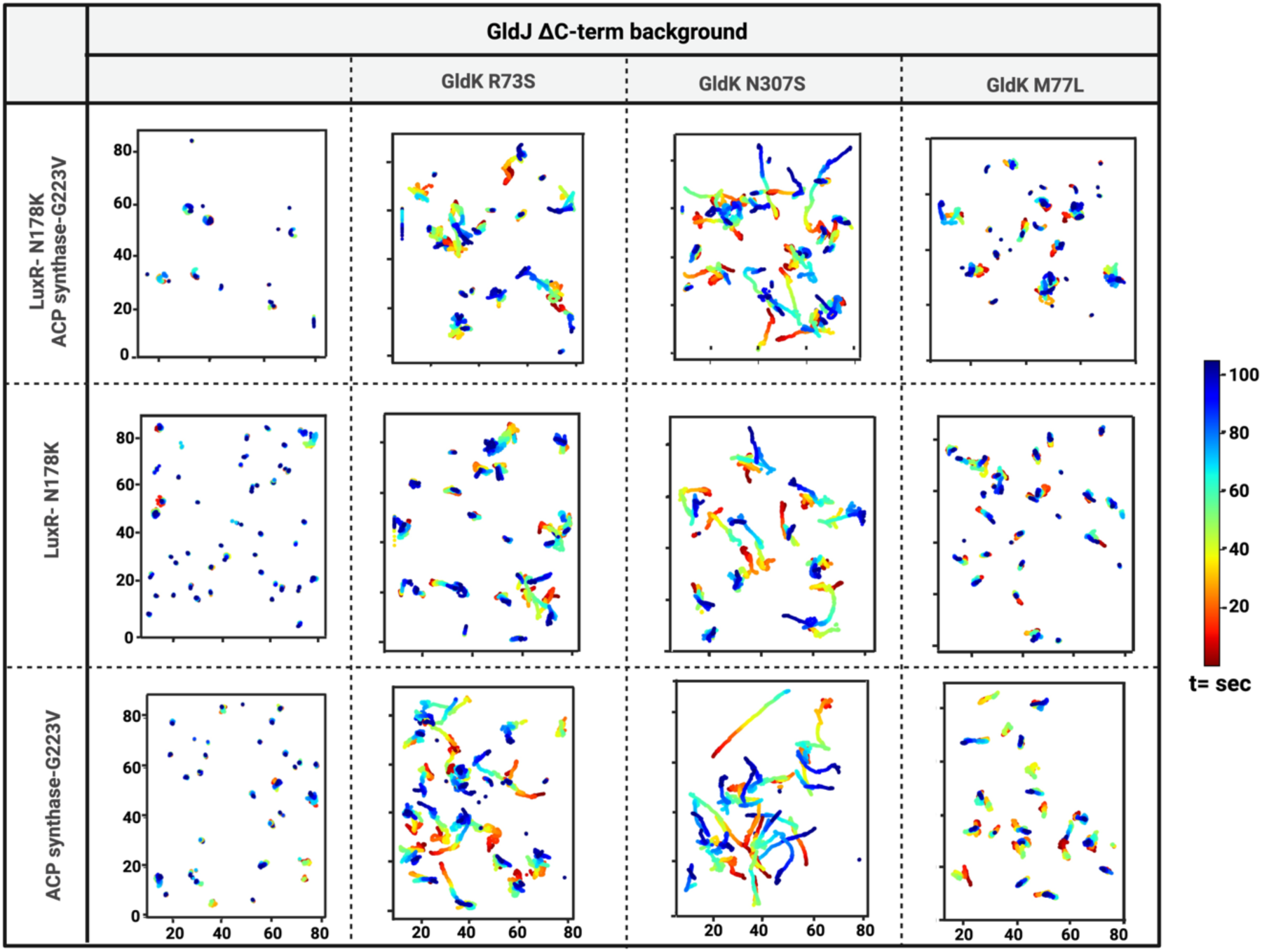
Microscopic imaging and single cell tracking over a glass surface provided further evidence that G223V mutation in ACP synthase and N178K mutation in a LuxR-like protein do not play a role in the restoration of motility.

**Figure S4.**
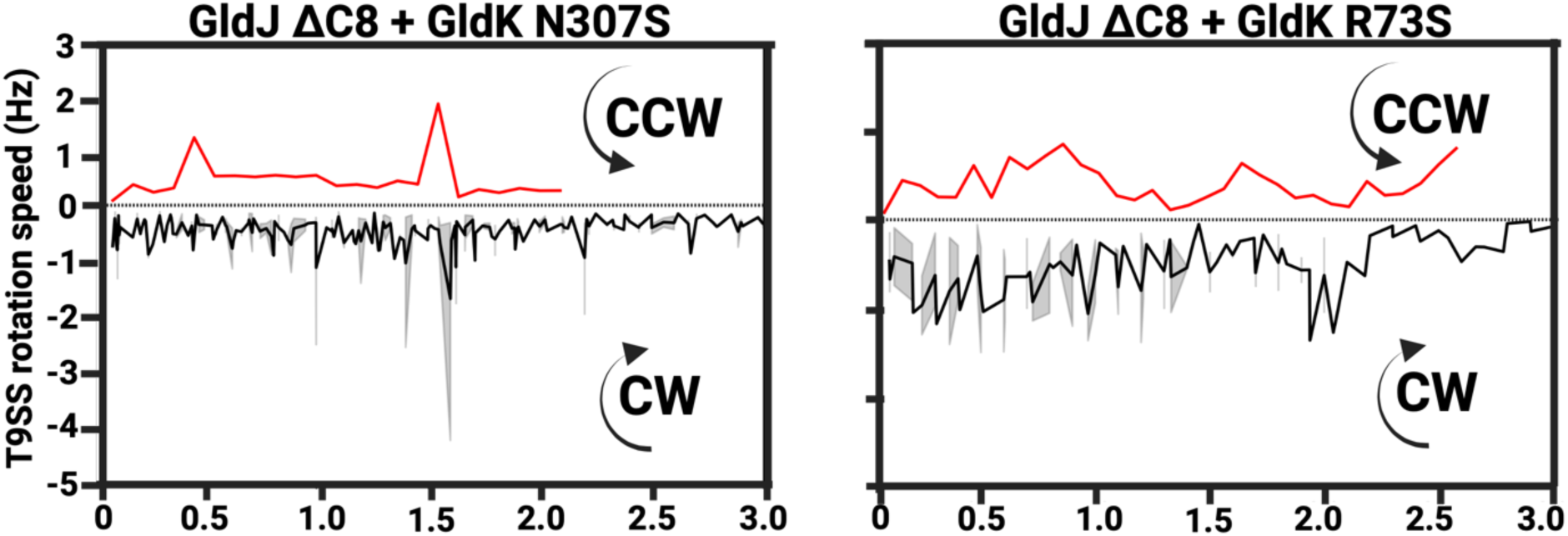
The rotational direction of T9SS motors of GldJ ΔC8 cells with GldK N307S mutation (evolved strain) is predominantly CW but unlike the parent GldJ ΔC8 cells where all T9SS motors are CW, few CCW motors (red) are also observed in the evolved strain.

**Figure S5.**
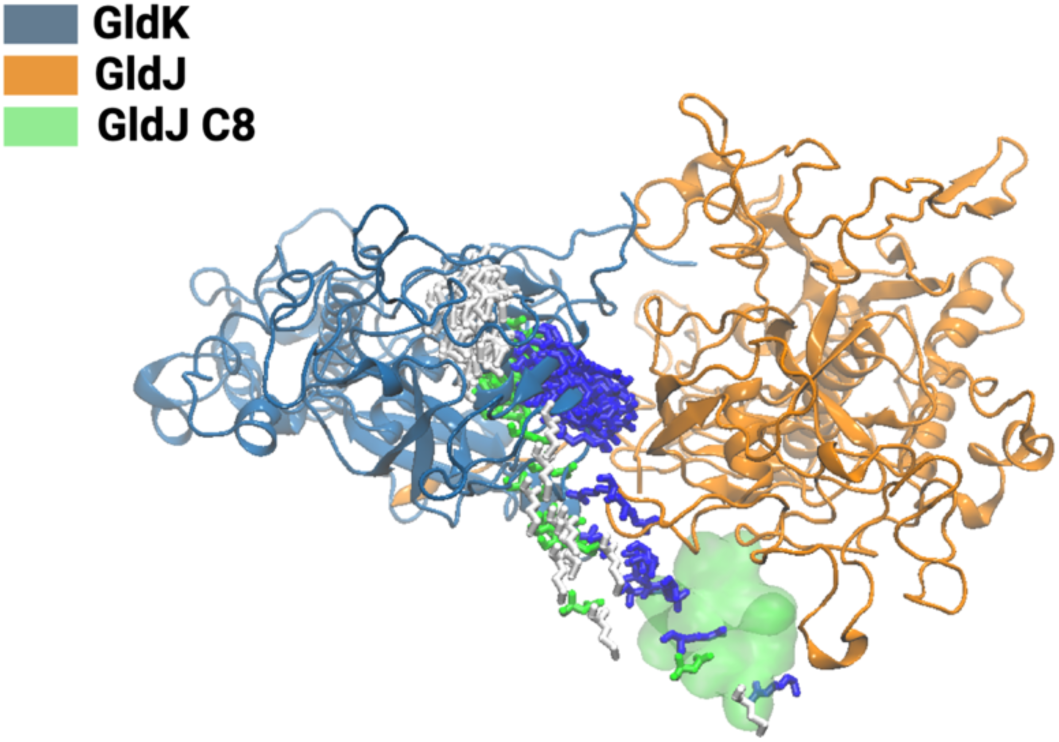
MD stimulations show that in the GldJ ΔC8 strain, the functional GldK residues identified by the suppressor screen (blue, white, and green stick diagrams) can dynamically replace the region previously occupied C8 of wild-type GldJ (shown as a green surface).

**Figure S6.**
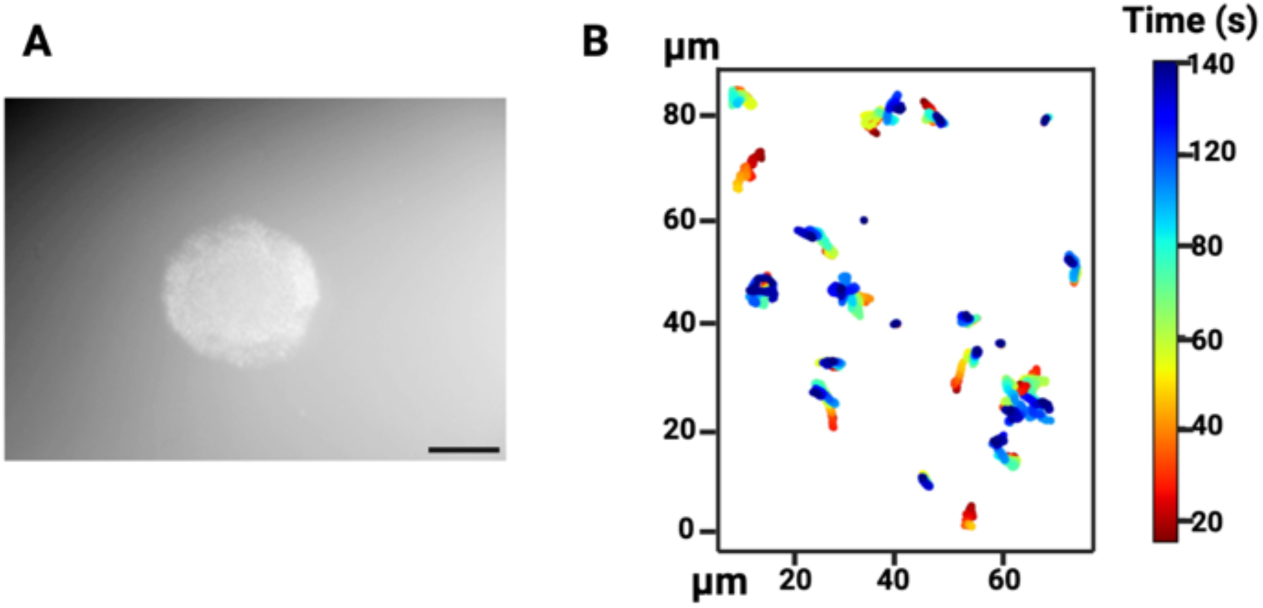
Combination of all three GldK point mutations (GldK R73S, M77L, and N307S) in a GldJ ΔC8 background does not restore (A) swarming on agar and (B) Single cell-motility over a glass surface. Scale bar for panel A is 6 mm.

**Figure S7.**
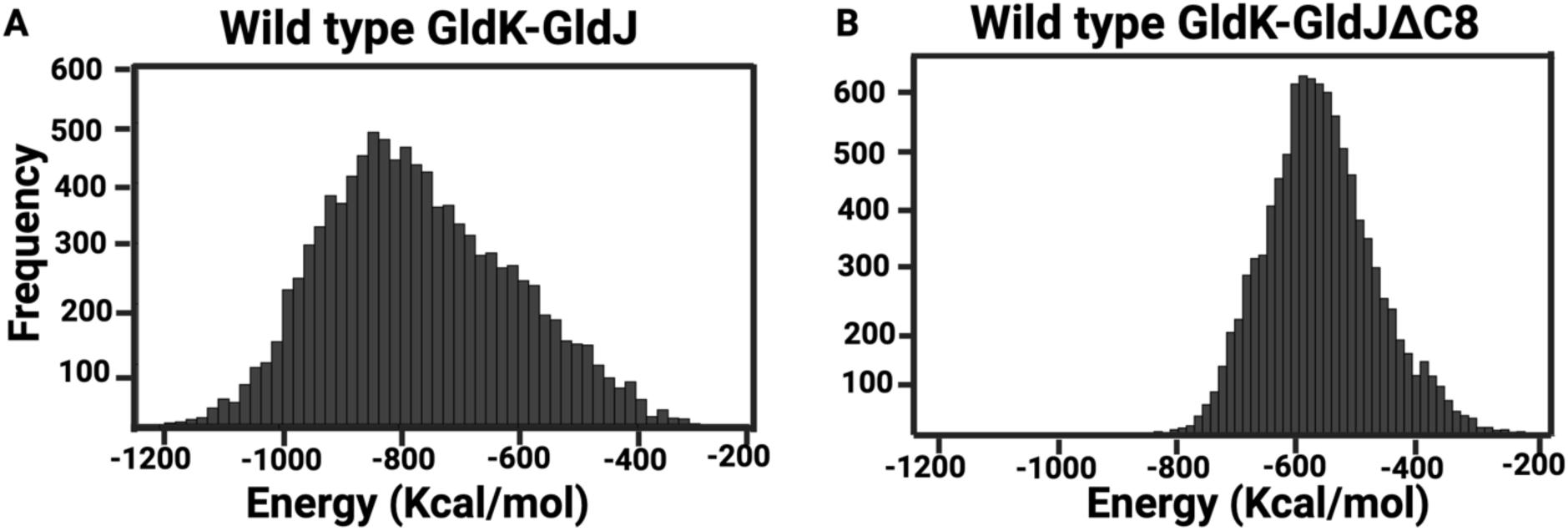
(A) Nonbonded interaction energy analysis illustrating a stronger interaction between GldJ and GldK in the wild-type GldJ and GldK system (−776.4 Kcal/mol average) compared to, (B) a weaker interaction in the GldJΔC8 and GldK system (−537.3 Kcal/mol average).

**Figure S8.**
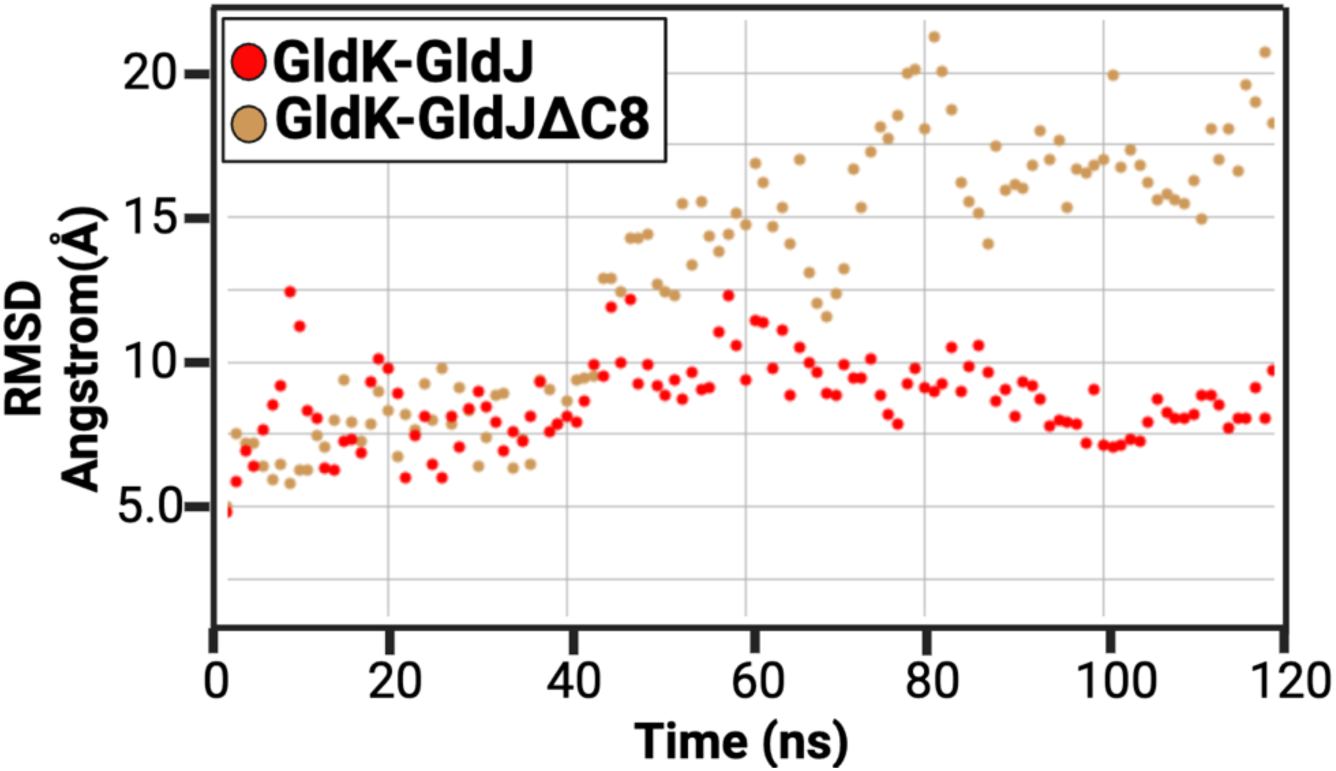
RMSD analysis of GldJ and GldK in both the WT and mutant systems. GldK deviates significantly from its initial structure in the GldJ ΔC8 model reaching a final RMSD of ∼20 Å, which is around 10 Å larger than the wild-type model.

**Figure S9.**
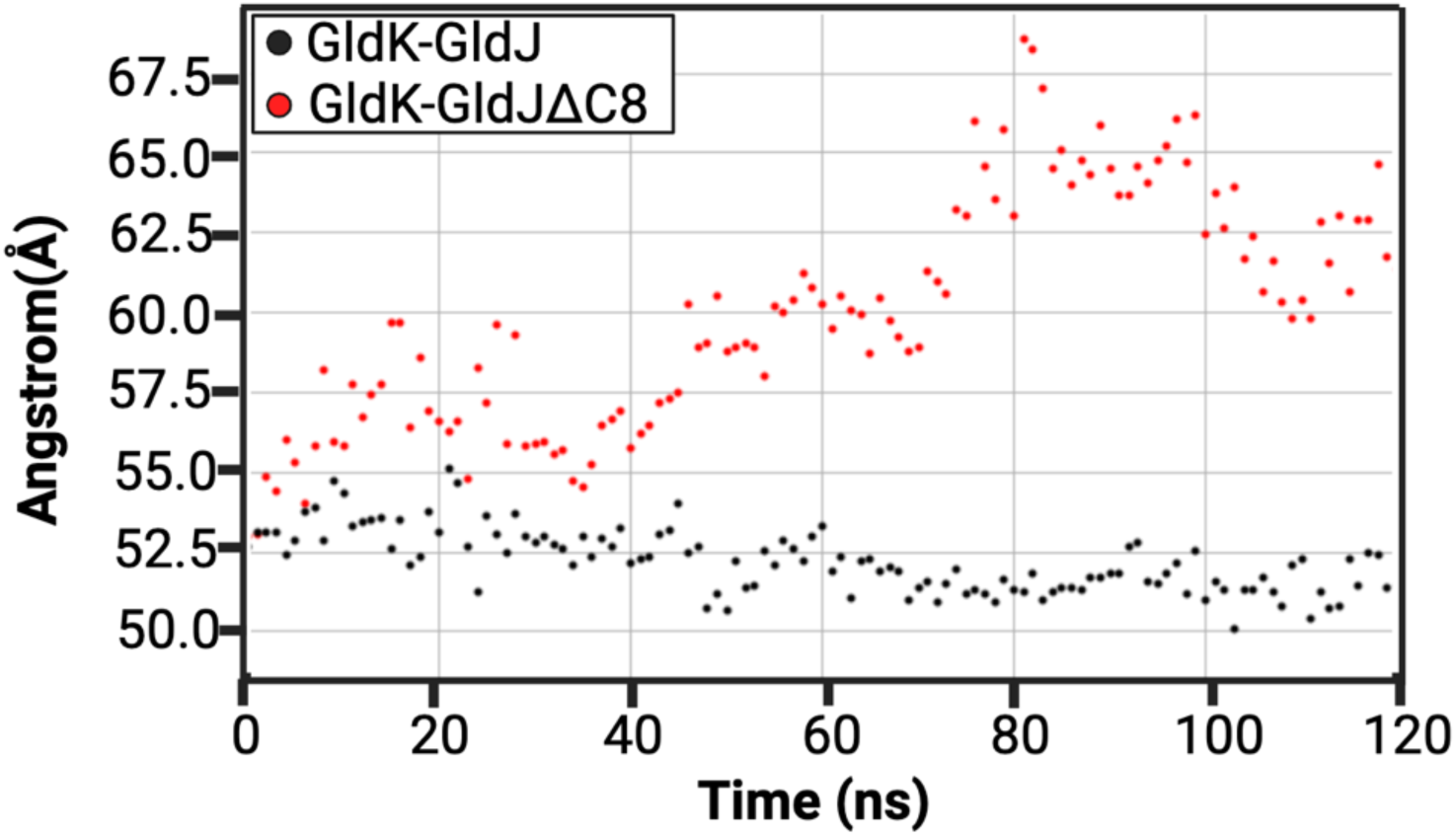
The inter center of mass distances, including all heavy atoms, between GldK and GldJ in the wild-type and GldJ ΔC8 systems recorded periodically throughout the MD simulation.

**Figure S10.**
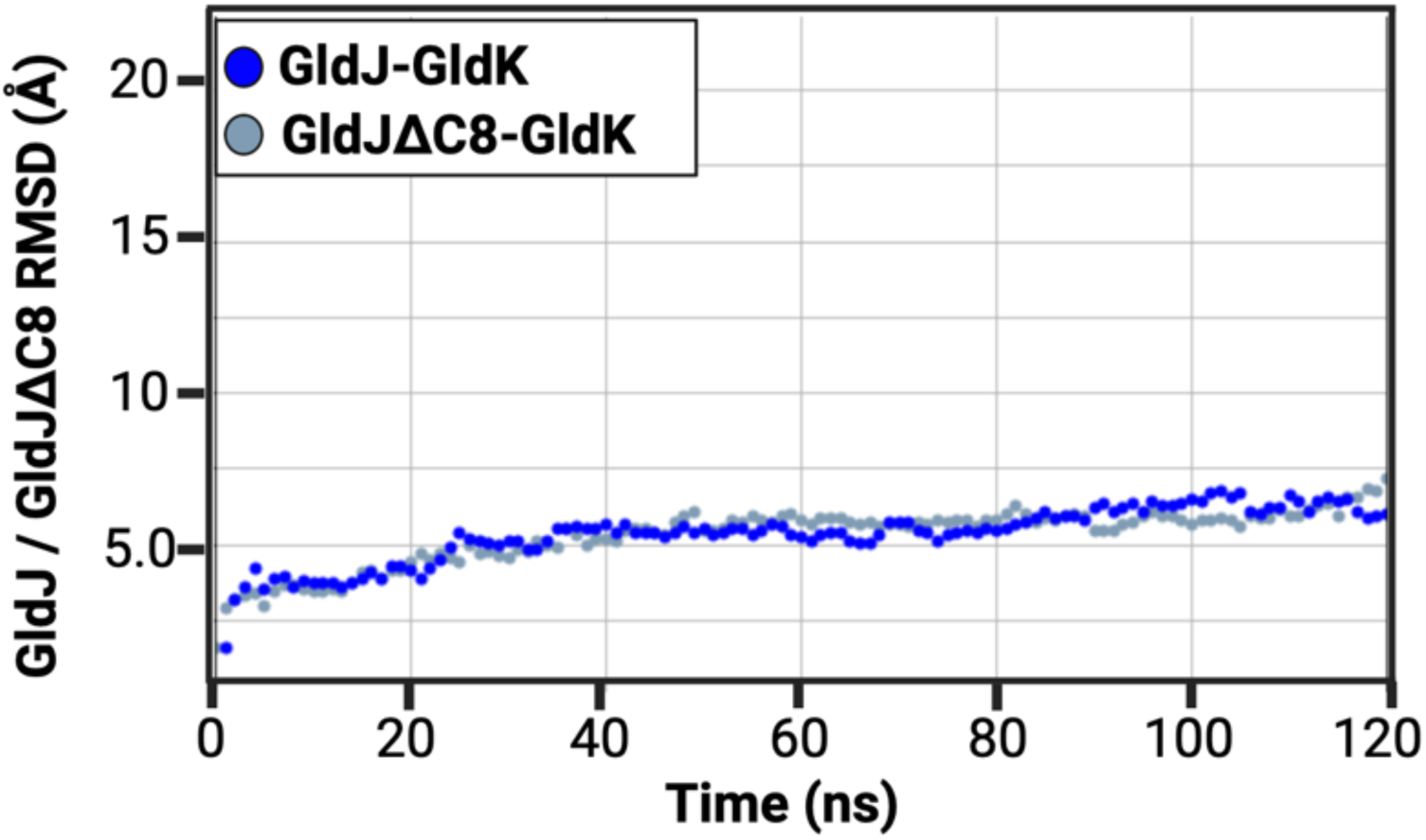
The RMSD of GldJ in the GldJ-GldK system, compared to GldJΔC8 in the GldJΔC8-GldK system, indicates that GldJ does not expand in a manner similar to GldK when C8 of GldJ is deleted.

**Figure S11.**
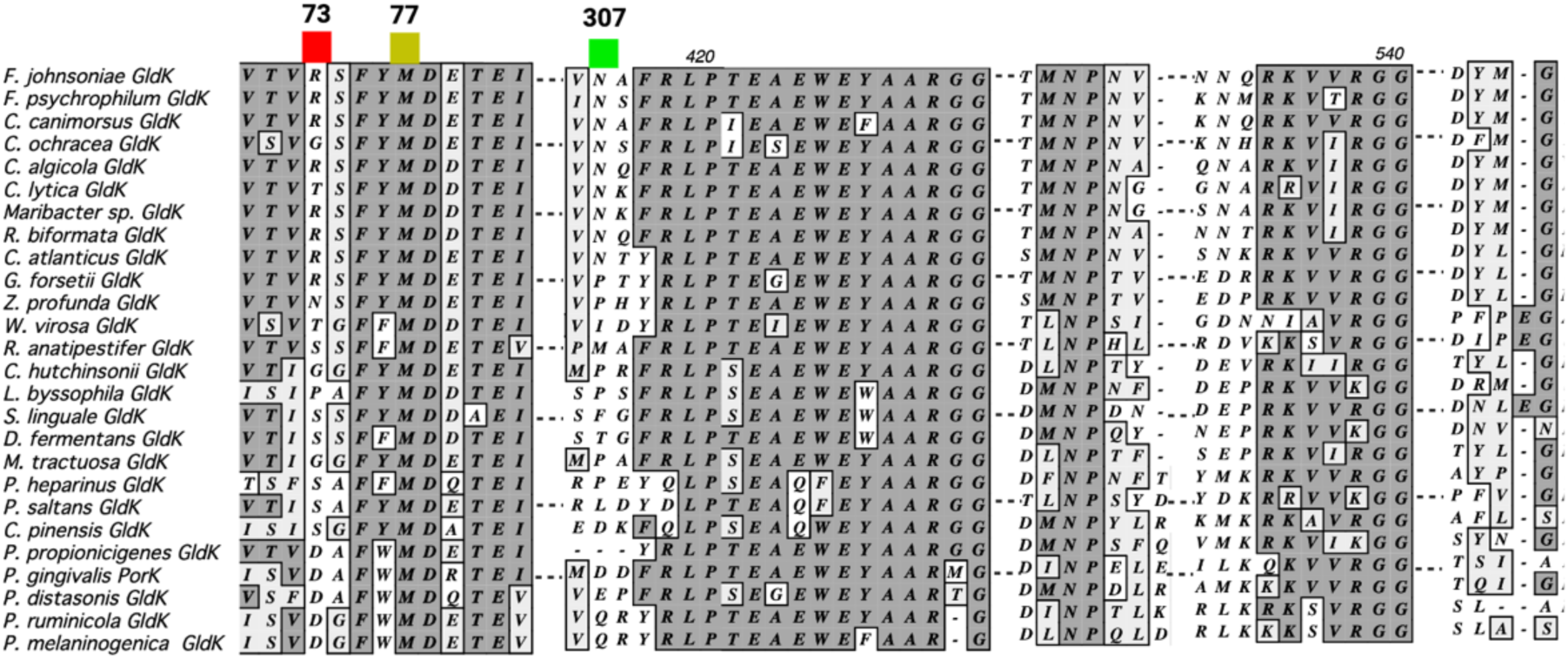
Multiple sequence alignment (MSA) of GldK protein, highlighting point mutations in moderately conserved regions across various species. Each mutation is indicated by a specific color, Red GldKR73S, Yellow GldK M77L and Green GldKN307S reflecting its evolutionary conservation and potential impact on protein function.

## SUPPLEMENTARY TEXT

### Single amino acid substitutions of LuxR and Acyl Carrier Protein (ACP) Synthase in a GldJ ΔC8 background does not improve motility

The three GldK suppressor strains isolated from our screen also had a N178K mutation in a putative LuxR-like protein (*Fjoh_4220*) and a G223V mutation in an Acyl Carrier Protein (ACP) Synthase (*Fjoh_3810*). No reports suggest that these two proteins influence Bacteroidetes gliding motility, and it’s plausible that these mutations were already present in the parent strain prior to the suppressor screen. To gain a deeper understand of their role, individual strains that carry LuxR N178K, ACP G223V, and LuxR N178K plus ACP G223V mutation in a GldJ ΔC-terminal background were generated. When comparing the new strains with mutations in LuxR and ACP to the original GldJ ΔC-terminal strain, there were no noticeable improvements in swarming and single-cell motility (Supplementary video 6, Supplementary video 7) The strains that had both GldK R73S, M77L, and N307S mutations, along with the LuxR and ACP mutations, showed similar swarming and single-cell motility characteristics as the strains with only GldK R73S, M77L, and N307S mutations in a GldJ ΔC-terminal background. Similarly, for LuxR N178K and ACP G223V individually GldJ ΔC-terminal background exhibited no motility. Furthermore, when both LuxR N178K and ACP G223V were present in individual GldK point mutants no improvement in speed was observed (Supplementary video 8, 9). These results indicate that the suppressor phenotype is not influenced by the LuxR N178K and ACP G223V mutations (Figs. S3, S4).

### Combinatorial mutagenesis of GldK point mutations in a GldJ ΔC-terminal background does not restore motility

A triple GldK mutant strain, which includes the mutations GldK R73S, M77L, and N307S in the GldJ ΔC8 background, was created. However, this mutant strain did not exhibit swarming on agar, and its single cells displayed sluggish back and forth motion (Fig. S6). These characteristics were comparable to the behavior observed in cells of the original GldJ ΔC8 strain.

## SUPPLEMENTARY TABLES

**Table S1.**
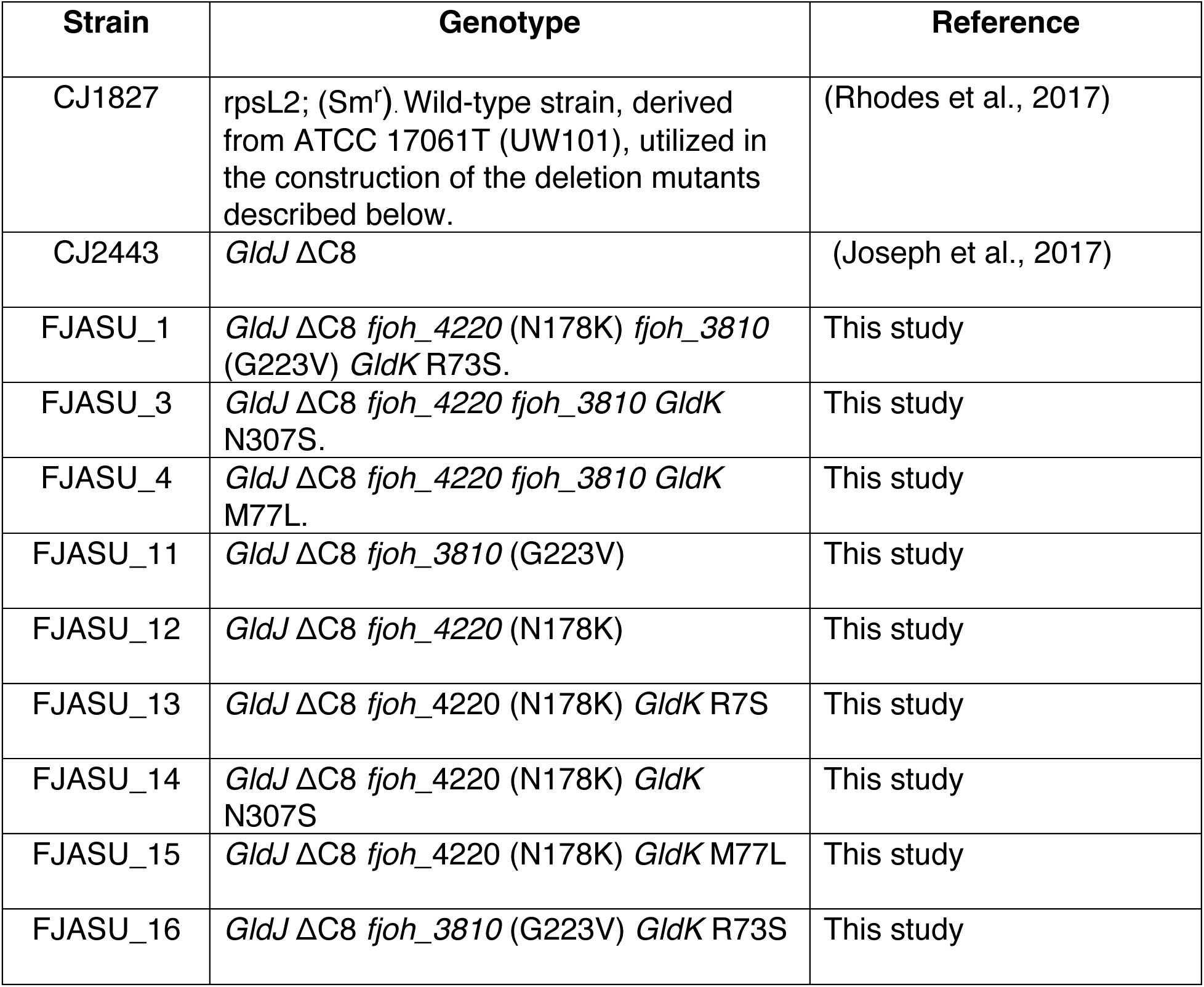
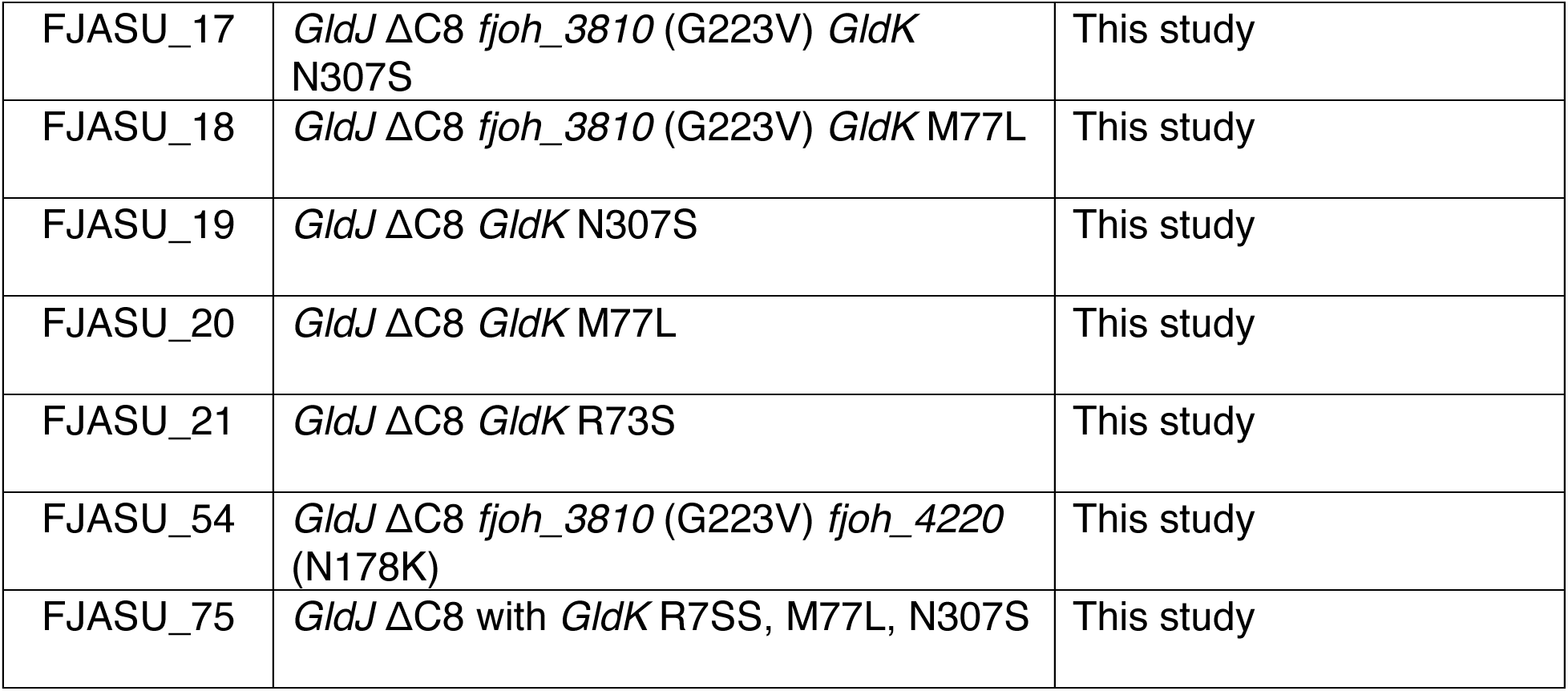
Strain List

| Strain | Genotype | Reference |
| --- | --- | --- |
| CJ1827 | rpsL2; (Sm<sup>r</sup>). Wild-type strain, derived from ATCC 17061T (UW101), utilized in the construction of the deletion mutants described below. | (Rhodes et al., 2017) |
| CJ2443 | *GldJ* ΔC8 | (Joseph et al., 2017) |
| FJASU_1 | *GldJ* ΔC8 *fjoh*_4220 (N178K) *fjoh*_3810 (G223V) *GldK* R73S. | This study |
| FJASU_3 | *GldJ* ΔC8 *fjoh*_4220 *fjoh*_3810 *GldK* N307S. | This study |
| FJASU_4 | *GldJ* ΔC8 *fjoh*_4220 *fjoh*_3810 *GldK* M77L. | This study |
| FJASU_11 | *GldJ* ΔC8 *fjoh*_3810 (G223V) | This study |
| FJASU_12 | *GldJ* ΔC8 *fjoh*_4220 (N178K) | This study |
| FJASU_13 | *GldJ* ΔC8 *fjoh*_4220 (N178K) *GldK* R7S | This study |
| FJASU_14 | *GldJ* ΔC8 *fjoh*_4220 (N178K) *GldK* N307S | This study |
| FJASU_15 | *GldJ* ΔC8 *fjoh*_4220 (N178K) *GldK* M77L | This study |
| FJASU_16 | *GldJ* ΔC8 *fjoh*_3810 (G223V) *GldK* R73S | This study |
| FJASU_17 | *GldJ* ΔC8 *fjoh_3810* (G223V) *GldK* N307S | This study |
| FJASU_18 | *GldJ* ΔC8 *fjoh_3810* (G223V) *GldK* M77L | This study |
| FJASU_19 | *GldJ* ΔC8 *GldK* N307S | This study |
| FJASU_20 | *GldJ* ΔC8 *GldK* M77L | This study |
| FJASU_21 | *GldJ* ΔC8 *GldK* R73S | This study |
| FJASU_54 | *GldJ* ΔC8 *fjoh_3810* (G223V) *fjoh_4220* (N178K) | This study |
| FJASU_75 | *GldJ* ΔC8 with *GldK* R73S, M77L, N307S | This study |

**Table S2.**
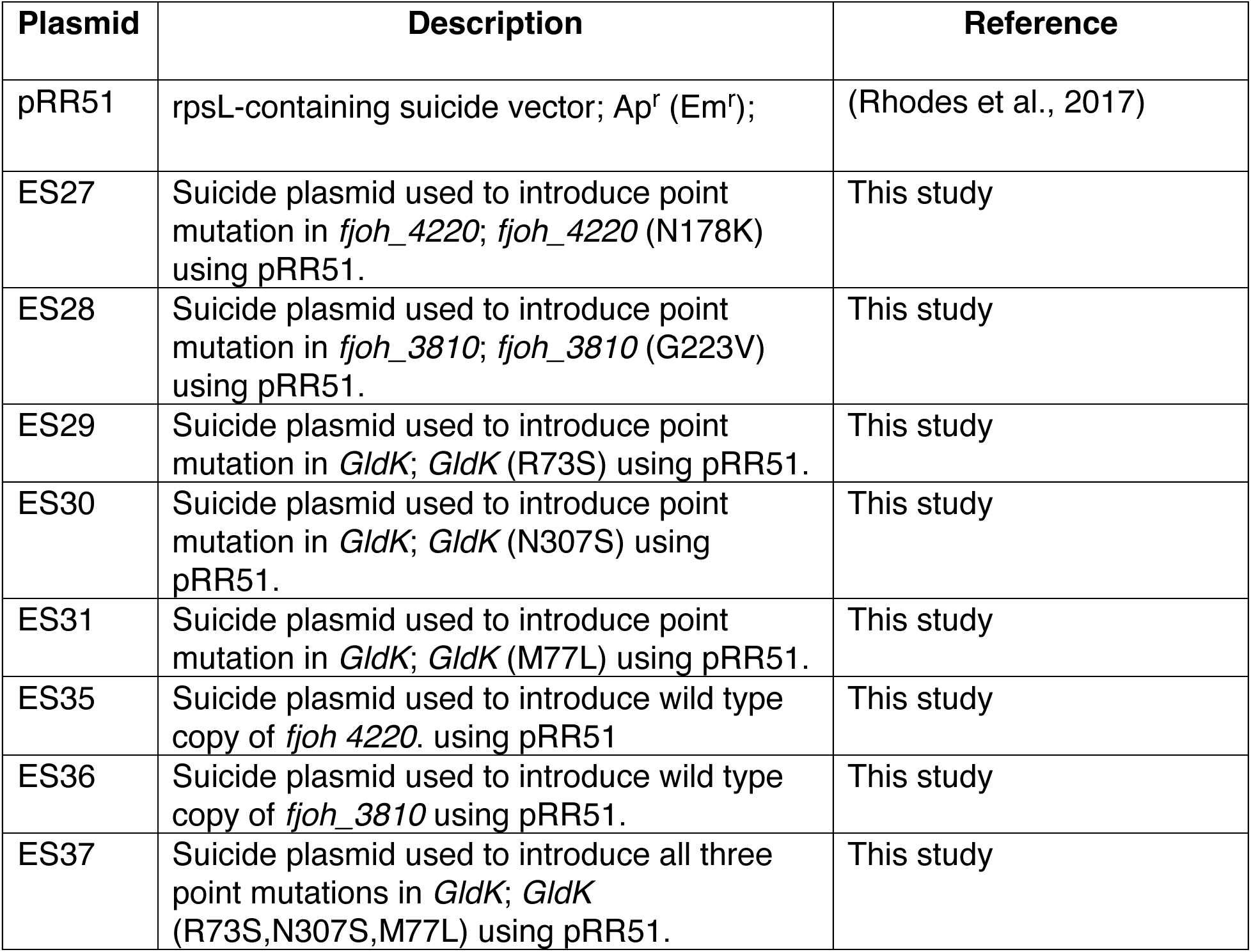
List of plasmids used in this study.

| Plasmid | Description | Reference |
| --- | --- | --- |
| pRR51 | rpsL-containing suicide vector; Ap<sup>r</sup> (Em<sup>r</sup>); | (Rhodes et al., 2017) |
| ES27 | Suicide plasmid used to introduce point mutation in *fjoh_4220*; *fjoh_4220* (N178K) using pRR51. | This study |
| ES28 | Suicide plasmid used to introduce point mutation in *fjoh_3810*; *fjoh_3810* (G223V) using pRR51. | This study |
| ES29 | Suicide plasmid used to introduce point mutation in *GldK*; *GldK* (R73S) using pRR51. | This study |
| ES30 | Suicide plasmid used to introduce point mutation in *GldK*; *GldK* (N307S) using pRR51. | This study |
| ES31 | Suicide plasmid used to introduce point mutation in *GldK*; *GldK* (M77L) using pRR51. | This study |
| ES35 | Suicide plasmid used to introduce wild type copy of *fjoh_4220*. using pRR51 | This study |
| ES36 | Suicide plasmid used to introduce wild type copy of *fjoh_3810* using pRR51. | This study |
| ES37 | Suicide plasmid used to introduce all three point mutations in *GldK*; *GldK* (R73S,N307S,M77L) using pRR51. | This study |

**Table S3.**
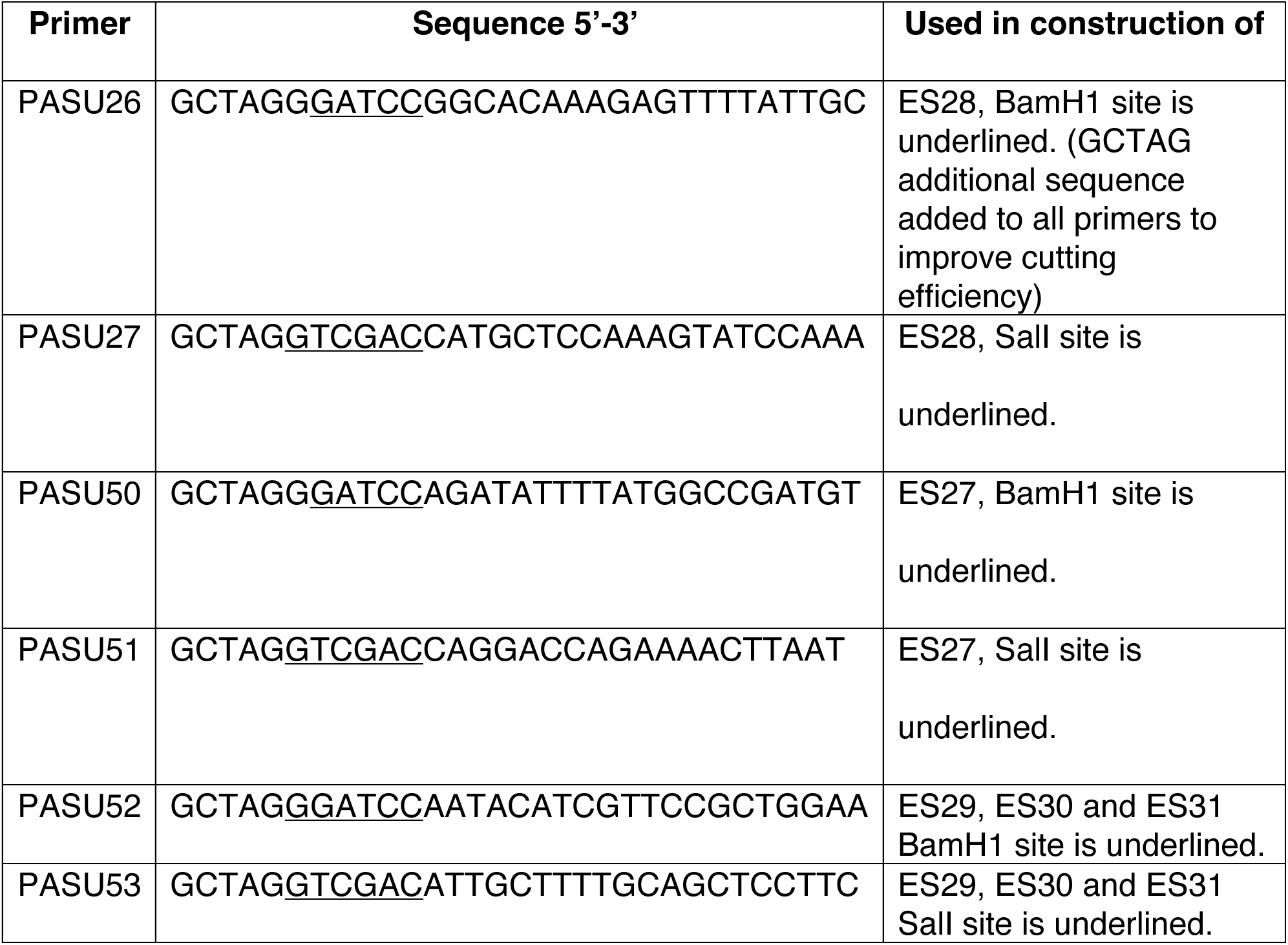
List of primers used in this study.

## SUPPLEMENTARY VIDEOS

**Supplementary video 1.** Timelapse images of a tethered wild-type cell with a counterclockwise rotating T9SS.

**Supplementary video 2.** Timelapse images of a tethered GldJ ΔC8 cell with a clockwise rotating T9SS.

**Supplementary video 3.** Timelapse images showing gliding of wild-type cells and GldJ ΔC8 cells over a glass surface.

**Supplementary video 4.** Timelapse images of cells containing GldK R73S, GldK M77L and GldK N307S individual point mutations in GldJ ΔC8 background over a glass surface.

**Supplementary video 5.** Timelapse TIRF images of immunofluorescent SprB motion on the surface of a wild-type cell, GldJ ΔC8 cell, GldK N307S in GldJ ΔC8 background.

**Supplementary video 6.** Timelapse images, over a glass surface, of GldJ ΔC8 cells containing GldK R73S ACP G223V, GldK M77L ACP G223V, and GldKN307S ACP G223V.

**Supplementary video 7.** Timelapse images, over a glass surface, of GldJ ΔC8 cells containing GldK R73S LuxR N178K, GldK M77L LuxR N178K, and GldKN307S LuxR N178K.

**Supplementary video 8.** Timelapse images, over a glass surface, of cells containing ACP_G223V and LuxR_N178K mutation individually a GldJ ΔC8 background and ACP_G223V and LuxR_N178K together in a GldJ ΔC8 background.

**Supplementary video 9.** Timelapse images, over a glass surface, of cells containing (i) GldK R73S, ACPG223V, and LuxR N178K mutations in a GldJ ΔC8 background. (ii) GldK R73S, ACP G223V, and LuxR N178K mutations in a GldJ ΔC8 background. (iii) GldK N307S, ACP G223V, and LuxR N178K mutations in a GldJ ΔC8 background.

**Supplementary video 10.** Time-lapse images from tethered cell assays of the evolved GldK mutant strains in a GldJ ΔC8 background, illustrating examples with both CW and CCW rotating T9SS.

## References.

1. Paysan-Lafosse, T. et al. InterPro in 2022. Nucleic Acids Research 51, D418–D427 (2023).

2. Shrivastava, A., Johnston, J. J., van Baaren, J. M. & McBride, M. J. Flavobacterium johnsoniae GldK, GldL, GldM, and SprA are required for secretion of the cell surface gliding motility adhesins SprB and RemA. J Bacteriol 195, 3201–3212 (2013).

3. Shrivastava, A., Lele, P. P. & Berg, H. C. A rotary motor drives Flavobacterium gliding. Curr Biol 25, 338–341 (2015).

4. Saiki, K. & Konishi, K. Identification of a Porphyromonas gingivalis novel protein sov required for the secretion of gingipains. Microbiol Immunol 51, 483–491 (2007).

5. Kharade, S. S. & McBride, M. J. Flavobacterium johnsoniae PorV is required for secretion of a subset of proteins targeted to the type IX secretion system. J Bacteriol 197, 147–158 (2015).

6. Zhu, Y. & McBride, M. J. The unusual cellulose utilization system of the aerobic soil bacterium Cytophaga hutchinsonii. Appl Microbiol Biotechnol 101, 7113–7127 (2017).

7. Li, Y., et al. Identification of trypsin-degrading commensals in the large intestine. Nature 609, 582–589 (2022).

8. Sato, K., et al. A protein secretion system linked to bacteroidete gliding motility and pathogenesis. Proc Natl Acad Sci U S A 107, 276–281 (2010).

9. Dominy, S. S., et al. Porphyromonas gingivalis in Alzheimer’s disease brains: Evidence for disease causation and treatment with small-molecule inhibitors. Sci Adv 5, eaau3333 (2019).

10. Kita, D., et al. Involvement of the Type IX Secretion System in Capnocytophaga ochracea Gliding Motility and Biofilm Formation. Appl Environ Microbiol 82, 1756–1766 (2016).

11. Shrivastava, A., et al. Cargo transport shapes the spatial organization of a microbial community. Proc Natl Acad Sci U S A 115, 8633–8638 (2018).

12. Ratheesh, N. K., Calderon, C. A., Zdimal, A. M. & Shrivastava, A. Bacterial Swarm-Mediated Phage Transportation Disrupts a Biofilm Inherently Protected from Phage Penetration. http://biorxiv.org/lookup/doi/10.1101/2021.06.25.449910 (2021) doi:10.1101/2021.06.25.449910.

13. Guo, Y., et al. Riemerella anatipestifer Type IX Secretion System Is Required for Virulence and Gelatinase Secretion. Front Microbiol 8, 2553 (2017).

14. Pisarenko, O., Studneva, I. & Khlopkov, V. Metabolism of the tricarboxylic acid cycle intermediates and related amino acids in ischemic guinea pig heart. Biomed Biochim Acta 46, S568–571 (1987).

15. Barbier, P., et al. The Type IX Secretion System Is Required for Virulence of the Fish Pathogen Flavobacterium psychrophilum. Appl Environ Microbiol 86, e00799–20 (2020).

16. How, K. Y., Song, K. P. & Chan, K. G. Porphyromonas gingivalis: An Overview of Periodontopathic Pathogen below the Gum Line. Front Microbiol 7, 53 (2016).

17. LaFrentz, B. R., et al. The fish pathogen Flavobacterium columnare represents four distinct species: Flavobacterium columnare, Flavobacterium covae sp. nov., Flavobacterium davisii sp. nov. and Flavobacterium oreochromis sp. nov., and emended description of Flavobacterium columnare. Systematic and Applied Microbiology 45, 126293 (2022).

18. Yin, C., et al. Rhizosphere community selection reveals bacteria associated with reduced root disease. Microbiome 9, 86 (2021).

19. Niraula, S., Rose, M. & Chang, W.-S. Microbial co-occurrence network in the rhizosphere microbiome: its association with physicochemical properties and soybean yield at a regional scale. J Microbiol 60, 986–997 (2022).

20. McBride, M. J. *Bacteroidetes* Gliding Motility and the Type IX Secretion System. Microbiol Spectr 7, 7.1.15 (2019).

21. Nakane, D., Sato, K., Wada, H., McBride, M. J. & Nakayama, K. Helical flow of surface protein required for bacterial gliding motility. Proceedings of the National Academy of Sciences 110, 11145–11150 (2013).

22. Shrivastava, A., Roland, T. & Berg, H. C. The Screw-Like Movement of a Gliding Bacterium Is Powered by Spiral Motion of Cell-Surface Adhesins. Biophysical Journal 111, 1008–1013 (2016).

23. Shibata, S., et al. Filamentous structures in the cell envelope are associated with bacteroidetes gliding machinery. Commun Biol 6, 94 (2023).

24. Trivedi, A., Gosai, J., Nakane, D. & Shrivastava, A. Design Principles of the Rotary Type 9 Secretion System. Front Microbiol 13, 845563 (2022).

25. Hennell James, R., et al. Structure and mechanism of the proton-driven motor that powers type 9 secretion and gliding motility. Nat Microbiol 6, 221–233 (2021).

26. Vincent, M. S., et al. *Dynamic Proton-Dependent Motors Power Type IX Secretion and Gliding Adhesin Movement in* Flavobacterium. http://biorxiv.org/lookup/doi/10.1101/2021.10.19.464928 (2021) doi:10.1101/2021.10.19.464928.

27. Song, L., et al. A unique bacterial secretion machinery with multiple secretion centers. Proc Natl Acad Sci U S A 119, e2119907119 (2022).

28. Sarkar, M. K., Paul, K. & Blair, D. Chemotaxis signaling protein CheY binds to the rotor protein FliN to control the direction of flagellar rotation in Escherichia coli. Proc Natl Acad Sci U S A 107, 9370–9375 (2010).

29. Chang, Y., et al. Molecular mechanism for rotational switching of the bacterial flagellar motor. Nat Struct Mol Biol 27, 1041–1047 (2020).

30. Glaser, J. & Pate, J. L. Isolation and characterization of gliding motility mutants of Cytophaga columnaris. Archiv. Mikrobiol. 93, 295–309 (1973).

31. Lapidus, I. R. & Berg, H. C. Gliding motility of Cytophaga sp. strain U67. J Bacteriol 151, 384–398 (1982).

32. Braun, T. F. & McBride, M. J. Flavobacterium johnsoniae GldJ is a lipoprotein that is required for gliding motility. J Bacteriol 187, 2628–2637 (2005).

33. Johnston, J. J., Shrivastava, A. & McBride, M. J. Untangling Flavobacterium johnsoniae Gliding Motility and Protein Secretion. J Bacteriol 200, e00362–17 (2018).

34. Shrivastava, A. & Berg, H. C. A molecular rack and pinion actuates a cell-surface adhesin and enables bacterial gliding motility. Sci Adv 6, eaay6616 (2020).

35. Kharade, S. S. & McBride, M. J. Flavobacterium johnsoniae chitinase ChiA is required for chitin utilization and is secreted by the type IX secretion system. J Bacteriol 196, 961–970 (2014).

36. Zhu, Y. & McBride, M. J. Deletion of the Cytophaga hutchinsonii type IX secretion system gene sprP results in defects in gliding motility and cellulose utilization. Appl Microbiol Biotechnol 98, 763–775 (2014).

37. Yan, Y., Tao, H., He, J. & Huang, S.-Y. The HDOCK server for integrated protein-protein docking. Nat Protoc 15, 1829–1852 (2020).

38. Vincent, M. S., et al. Characterization of the Porphyromonas gingivalis Type IX Secretion Trans-envelope PorKLMNP Core Complex. J Biol Chem 292, 3252–3261 (2017).

39. Rieu, M., Krutyholowa, R., Taylor, N. M. I. & Berry, R. M. A new class of biological ion-driven rotary molecular motors with 5:2 symmetry. Front Microbiol 13, 948383 (2022).

40. Shekhar, M., et al. CryoFold: determining protein structures and data-guided ensembles from cryo-EM density maps. Matter 4, 3195–3216 (2021).

41. Gorasia, D., et al. In Situ *Structure and Organisation of the Type IX Secretion System*. http://biorxiv.org/lookup/doi/10.1101/2020.05.13.094771 (2020) doi:10.1101/2020.05.13.094771.

42. Gorasia, D. G., et al. Protein Interactome Analysis of the Type IX Secretion System Identifies PorW as the Missing Link between the PorK/N Ring Complex and the Sov Translocon. Microbiol Spectr e01602–21 (2022) doi:10.1128/spectrum.01602-21.

43. Chang, Y., Carroll, B. L. & Liu, J. Structural basis of bacterial flagellar motor rotation and switching. Trends in Microbiology 29, 1024–1033 (2021).

44. Gorasia, D. G., et al. Structural Insights into the PorK and PorN Components of the Porphyromonas gingivalis Type IX Secretion System. PLoS Pathog 12, e1005820 (2016).

45. El-sayed, M. E., Youssef, A. W., Shehata, O. M., Shihata, L. A. & Azab, E. Computer vision for package tracking on omnidirectional wheeled conveyor: Case study. Engineering Applications of Artificial Intelligence 116, 105438 (2022).

46. Rhodes, R. G., Pucker, H. G. & McBride, M. J. Development and use of a gene deletion strategy for Flavobacterium johnsoniae to identify the redundant gliding motility genes remF, remG, remH, and remI. J Bacteriol 193, 2418–2428 (2011).

47. Nelson, S. S., Bollampalli, S. & McBride, M. J. SprB is a cell surface component of the Flavobacterium johnsoniae gliding motility machinery. J Bacteriol 190, 2851– 2857 (2008).

